# Parental origin of Gsα inactivation differentially affects bone remodeling in a mouse model of Albright hereditary osteodystrophy

**DOI:** 10.1101/2021.07.27.453811

**Authors:** Patrick McMullan, Peter Maye, Qingfen Yang, David W. Rowe, Emily L. Germain-Lee

**Affiliations:** Department of Pediatrics, University of Connecticut School of Medicine, Farmington, CT; Department of Reconstructive Sciences, University of Connecticut School of Dental Medicine, Farmington, CT; Center for Regenerative Medicine and Skeletal Development, University of Connecticut School of Dental Medicine, Farmington, CT; Albright Center, Division of Pediatric Endocrinology, Connecticut Children’s, Farmington, CT

**Keywords:** *Gnas*, bone, Genetic Animal Models, Molecular pathways – remodeling, Osteoblasts, Osteoclasts, Diseases and Disorders Of/Related to Bone – Other

## Abstract

Albright hereditary osteodystrophy (AHO) is caused by heterozygous inactivation of *GNAS*, a complex locus that encodes the alpha-stimulatory subunit of GPCRs (Gsα) in addition to *NESP55* and *XL*α*s* due to alternative first exons. AHO skeletal manifestations include brachydactyly, brachymetacarpia, compromised adult stature, and subcutaneous ossifications. AHO patients with maternally-inherited *GNAS* mutations develop pseudohypoparathyroidism type 1A (PHP1A) with resistance to multiple hormones that mediate their actions through GPCRs requiring Gsα (eg., PTH, TSH, GHRH, calcitonin) and severe obesity. Paternally-inherited *GNAS* mutations cause pseudopseudohypoparathyroidism (PPHP), in which patients have AHO skeletal features but do not develop hormonal resistance or marked obesity. These differences between PHP1A and PPHP are caused by tissue-specific reduction of paternal Gsα expression. Previous reports in mice have shown loss of *Gsα* causes osteopenia due to impaired osteoblast number and function and suggest AHO patients could display evidence of reduced bone mineral density (BMD). However, we previously demonstrated PHP1A patients display normal-increased BMD measurements without any correlation to body mass index or serum PTH. Due to these observed differences between PHP1A and PPHP, we utilized our laboratory’s AHO mouse model to address *whether Gsα* heterozygous inactivation by the targeted disruption of exon 1 of *Gnas* differentially affects bone remodeling based on the parental inheritance of the mutation. Mice with paternally-inherited (*GnasE1+/−p*) and maternally-inherited (*GnasE1+/−m*) mutations displayed reductions in osteoblasts along the bone surface compared to wildtype. *GnasE1+/−p* mice displayed reduced cortical and trabecular bone parameters due to impaired bone formation and excessive bone resorption. *GnasE1+/−m* mice however displayed enhanced bone parameters due to increased osteoblast activity and normal bone resorption. These distinctions in bone remodeling between *GnasE1+/−p* and *GnasE1+/−m* mice appear to be secondary to changes in the bone microenvironment driven by calcitonin-resistance within *GnasE1+/−m* osteoclasts and therefore warrant further studies into understanding how Gsα influences osteoblast-osteoclast coupling interactions.

## Introduction

Fuller Albright *et al.*^(1)^ identified a group of patients who presented with a ‘short, thickset figure,’ and biochemical abnormalities that included increased serum PTH levels, hypocalcemia and hyperphosphatemia. Further analysis revealed that these patients displayed a blunted physiologic response to PTH administration that included reduced urinary cAMP and phosphate excretion.^(1)^ Because these patients displayed typical biochemical features of hypoparathyroidism (low serum calcium with elevated serum phosphate levels) and similar patients were later shown to have elevated PTH levels due to end organ resistance, this disorder was termed “pseudohypoparathyoridism” (PHP).^(1)^ Future monitoring of similar patients by Albright *et al.*^(2)^ led to the identification of a patient who presented with the same physical features of PHP but displayed normal serum calcium, phosphate, and PTH levels. In addition, this patient exhibited a normal phosphaturic response to PTH administration; therefore, this patient was diagnosed with the disorder “pseudopseudohypoparathyroidism” (PPHP).^(2)^ Given that PHP and PPHP share a unique cluster of physical features, which include shortened adult stature, brachydactyly, brachymetacarpia, and the formation of subcutaneous ossifications, these skeletal manifestations are collectively referred to as Albright hereditary osteodystrophy (AHO) (for review 3-5).^(1–6)^

AHO is caused by the heterozygous inactivation of *GNAS,* the gene that encodes the α-stimulatory subunit (Gsα) of G protein-coupled receptors (GPCRs) and that is required for appropriate signal transduction of multiple hormones to activate adenylyl cyclase.^(3–5,7)^ The *GNAS* locus is very complex given that it encodes not only *Gsα* but also Neuroendocrine Secretory Protein 55 (*NESP55*) and Extra Large alpha-stimulatory subunit (*XL*αs) through the use of alternative first exons. ^(3–5,7–9)^ Furthermore, the *GNAS* locus is highly complex in that it is controlled by genomic imprinting, which can lead to partial transcriptional repression of one parental allele.^(3–5,7–9)^ Consequently, AHO patients can display a distinct subset of extraskeletal features dependent upon the inheritance pattern of their acquired mutation. Patients with a *Gsα* mutation on the maternally-inherited allele develop PHP type 1A (PHP1A) and display resistance to multiple hormones that mediate their actions through GPCRs requiring Gsα (e.g. PTH, TSH, GHRH, calcitonin, LH/FSH) and have severe obesity.^(5,10,19,11–18)^ Alternatively, patients with a *Gsα* mutation on the paternally-inherited allele develop PPHP and display AHO skeletal features without exhibiting hormonal resistance or severe obesity.^(20)^ The presence of hormonal resistance within PHP1A has been shown to be due to tissue-specific reduction of paternal *Gsα* expression within certain endocrine organs such as the pituitary^(21)^, thyroid,^(13,17,18)^ gonads^(16)^ and renal cortex.^(3,18,22,23)^ Based on mouse models, the severe obesity associated with PHP1A is most likely secondary to reduced paternal *Gsα* expression in the dorsomedial nucleus of the hypothalamus.^(24,25)^ It is critical, however, to acknowledge that genomic imprinting does not occur within all tissues. Rather, *Gsα* has been confirmed to be biallelically expressed in tissues and cell types such as the skin, heart, white adipocytes and chondrocytes.^(18,23,26–29)^ This biallelic expression in certain tissues and cell types is evident by the fact that PPHP and PHP1A patients share similar skeletal manifestations such as shortened adult stature and brachydactyly (which are proposed to be caused by accelerated chondrocyte differentiation due to impaired PTH/PTHrP signaling^(26,30,31)^) and subcutaneous ossification formation (which are potentially thought to originate from the accelerated osteogenic differentiation of cell types within the dermis and subcutaneous tissue).^(32–35)^

In the context of skeletal development, *Gsα* has been shown to serve an essential role in maintaining bone homeostasis by regulating osteoblast and osteocyte function.^(36–41)^ Conditional deletion of *Gsα* specifically within mesenchymal (*Prx1-CRE*^(37)^), osteoblast (both *Osx-CRE*^(39)^ *and Col1a-CRE*^(36)^) and osteocyte (*Dmp1-CRE*^(40,41)^) lineages led to a reduction in cortical and trabecular bone quality due to decreases in osteoblast number and function.^(36–41)^ Previous reports have suggested evidence of biallelic expression of *Gsα* from whole human skull and femur tissue samples^(27)^; however, no analysis of *Gsα* expression within specific cell types within the bone (osteoblast, osteoclast or osteocyte) have been evaluated to date. These findings would suggest that AHO patients could display evidence of reduced bone mineral density (BMD) and potentially be at an elevated risk of fracture. Paradoxically, however, we found that PHP1A patients display normal to increased BMD without any correlation to body mass index (BMI) or serum PTH measurements.^(42)^ These data suggest the enhanced BMD within PHP1A subjects may be attributed to underlying changes in osteoblast and/or osteoclast function during the bone remodeling process and that these changes could potentially be the result of *Gsα* imprinting within specific cell types in the bone microenvironment. In addition, early evidence in our mouse model suggested that these changes could be driven by underlying abnormalities in the bone remodeling process, specifically decreased osteoclast activity.^(18)^

Our AHO mouse model, generated by targeted disruption of exon 1 of *Gnas*^(18,43)^ *(Gnas E1+/−*) recapitulates many of the hormonal and metabolic features that are observed clinically among AHO patients; these features were shown to be due to *Gsα* imprinting.^(18)^ This model also recapitulates skeletal features of AHO including the formation of subcutaneous ossifications that occur due to *Gsα* haploinsufficiency.^(18,32)^ Initial skeletal characterization of this model did not identify significant changes in femur trabecular bone volume or osteoblast number of *Gnas E1+/−* mice compared to wild type (WT) at 5 months or 1 year of age.^(18)^ However, our initial studies of the skeletal characterization did not separate *Gnas E1+/−* mice based upon maternally-inherited (*Gnas E1+/−m*) or paternally-inherited (*Gnas E1+/−p*) mutations; additionally, these analyses did not evaluate the cortical bone quality of *Gnas E1+/−* or WT mice, which is a significant contributor to overall BMD. A subsequent study by Ramaswany *et al.*^(44)^ utilizing our AHO mouse model provided evidence of distinct changes in cortical bone architecture among adult *Gnas E1+/−p* and *Gnas E1+/−m* mice due to changes in endosteal osteoclast number. At 2 weeks of age, both *Gnas E1+/−m* and *Gnas E1+/−p* mice displayed reduced cortical bone thickness and BV/TV as measured by microcomputed tomography (microCT or MCT) compared to WT. MCT analysis at 3 and 9 months revealed *Gnas E1+/−p* mice maintained these reduced cortical bone parameters; however, *Gnas E1+/−m* mice displayed an elevated cortical bone thickness and BV/TV which were comparable to WT. These distinctions in cortical bone quality among *Gnas E1+/−* mice were found to correlate directly with changes in the number of osteoclasts on the endocortical surface; *Gnas E1+/−p* mice displayed an elevated number of endocortical osteoclasts, whereas *Gnas E1+/−m* displayed a similar number of endocortical osteoclasts compared to littermate controls.^(44)^ This observed inheritance-specific variation in osteoclast number between *Gnas E1+/−* mice *in vivo* warrants a deeper investigation into identifying the role of *Gsα* in regulating osteoblast-osteoclast coupling during the bone remodeling process.

The purpose of this work was to identify whether *Gsα* heterozygous inactivation can differentially affect osteoblast activity and bone formation based upon the inheritance pattern of the *Gnas* mutation. We provide evidence demonstrating that both *Gnas E1+/−m* and *Gnas E1+/− p* mice have a reduction in total osteoblasts on the bone surface when compared to WT. However, *Gnas E1+/−m* osteoblasts displayed substantially elevated bone formation activity when compared to both WT and *Gnas E1+/−p in vivo.* This elevated activity within *Gnas E1+/−m* mice directly correlated with evidence of elevated cortical and trabecular bone architecture when compared to *Gnas E1+/−p* mice. We also provide evidence that these observed changes in osteoblast activity *in vivo* are not driven by cell-autonomous changes in *Gnas E1+/−m* osteoblasts but rather may be secondary to partial resistance to calcitonin, a G protein-coupled hormone, within the osteoclast lineage, consistent with what has been previously observed in PHP1A patients (44, or for Review 5, 8, 44-46).^(5,8,45–47)^ These findings correlate with our previous clinical observations and are the first to provide direct evidence of *Gsα* differentially influencing bone remodeling by affecting osteoblast-osteoclast interactions based upon the inheritance pattern of the *Gsα* mutation. In addition, our results further implicate that calcitonin signaling within the osteoclast may serve a physiologic role in regulating bone formation.

## Materials and Methods

### Generation and maintenance of mice

The generation of mice carrying a targeted disruption of exon 1 of *Gnas (Gnas E1+/−)* was described previously.^(18,32)^ Mice were maintained on a pure 129SvEv background and were genotyped by PCR analysis. Each 20 μL reaction was performed with 2 μL DNA, 3μL of 10x Standard Taq Reaction Buffer (NEB), 0.1 μL Taq Polymerase (NEB), 40μM of *Gnas* Forward, *Gnas* Reverse and *Neo1* primers (sequences below), 10μM of DNTP mix (Promega), and 18 μL of InVitrogen Ultrapure Distilled Water. Reaction parameters were carried out as follows: 95°C for 5 minutes, 34 cycles of: (1) 95°C for 30 seconds; (2) 60°C for 30 seconds; (3) 72°C for 30 seconds, and 2 holds of 72°C for 5 minutes.

All mice that carry a mutant maternal *Gnas* allele are hereafter referred to as *Gnas E1+/− m* and those with a mutant paternal allele as *Gnas E1+/−p*. Wild type mice (*Gnas+/+*) are referred to as WT. Mice were fed a standard diet of Prolab RMH2500 mouse chow and water *ad libitum*. For each of the methods outlined, both male and female mice were utilized. All *in vitro* cell culture experiments were performed using both male and female 6-9 week old mice; however, all other experiments outlined were performed using 12 week old mice. All mouse protocols were carried out in accordance with the standards of the UConn Health Animal Care and Use Committee.

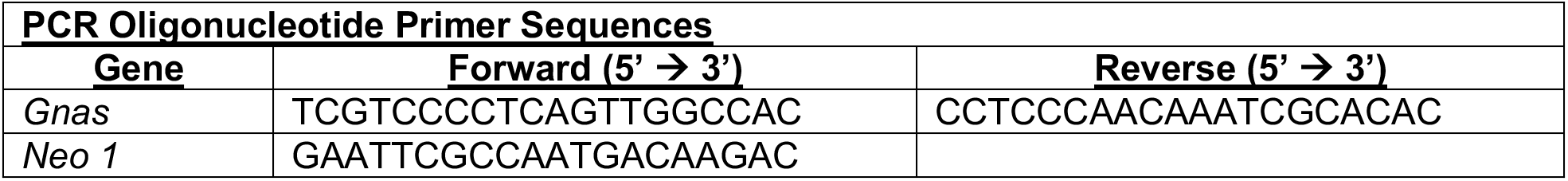

### Primary Bone Marrow Macrophage Cell Culture

The hind limbs of 6-9 week old WT and *Gnas E1+/−*mice were dissected using aseptic technique. After removing the epiphyses, the bone marrow was harvested from both femurs and tibias by flushing the marrow space with serum-free αMEM (Gibco) using a 26 gauge needle. The entire bone marrow harvested from one mouse was seeded onto a 10cm^2^ tissue culture-treated dish in αMEM with 10% Fetal Bovine Serum (FBS) and 1% Penicillin/Streptomycin. The following day, non-adherent cells were collected and underwent ficoll-hypaque gradient centrifugation to isolate bone marrow macrophages. To promote osteoclast differentiation, BMMs were cultured in αMEM with 10% FBS, 1% Penicillin/Streptomycin, as well as 30 ng/mL recombinant murine macrophage-colony stimulating factor (m-CSF, Peprotech) and 45 ng/mL recombinant murine RANK ligand (RANKL, Peprotech) for 5 days. For chromogenic TRAP staining, BMM cultures were seeded at 5×10^3^ cells/well of a 96-well dish. All additional *in vitro* BMM experiments were performed with 1×10^5^ cells/well in a 12-well dish. For staining analyses, BMM cultures were fixed with 5% glutaraldehyde in PBS for 10 minutes at room temperature, permeablized for 60 minutes at room temperature using 0.3% Triton-X and 0.1% BSA in PBS with calcium and magnesium. Cells were stained with TRITC-conjugated Phalloidin (Sigma Aldrich) for 45 minutes, followed by DAPI (Sigma Aldrich) for 10 minutes. BMM cultures were visualized by fluorescent microscopy utilizing both TRITC and DAPI filters. Following fluorescent image acquisition, BMM cultures were subsequently stained for chromogenic TRAP using a Leukocyte Acid Phosphatase Kit (Sigma 387A-1KT) for 60 minutes at 37°C as recommended by the manufacturer.

### Primary Bone Marrow Stromal Cell Culture

The hind limbs of 6-9 week old WT and *Gnas E1+/−* were dissected using aseptic technique. After removing the epiphyseal ends, the bone marrow was harvested from both femurs and tibias by flushing with a 26 gauge needle and the number of mononucleated cells were counted using a hemocytometer. For colony forming unit assays, 1×10^6^ mononucleated cells/well were seeded into a 6-well culture dish. For RNA analysis, as well as Alizarin Red and Von Kossa staining, 1×10^7^ cells/well were seeded into a 6-well dish. BMSCs were cultured for 7 days (Days -7 → 0) in basal culture media (αMEM with 10% FBS and 1% Penicillin/Streptomycin). Following this period, BMSCs underwent osteogenic differentiation by supplementing the basal media with 50μg/mL Ascorbic Acid and 10mM Beta-glycerophosphate for up to 14 days (Days 0 → 14).To monitor the number of osteoprogenitor colony formation units, alkaline phosphatase staining was completed within each well using a commercially available Leukocyte Alkaline Phosphatase staining kit (Sigma Aldrich 86R-1KT). Whole well images were collected through brightfield microscopy and quantification of colony forming units (CFUs) was performed using ImageJ. In order to monitor osteogenic differentiation of WT and *Gnas E1+/−*BMSCs, cultures were fixed for 10 minutes in 4% PFA in PBS for Alizarin Red Staining, or ice-cold 100% methanol for Von Kossa staining. Alizarin Red staining and quantification was performed using a commercially available staining kit (ScienCell Catalog #0223). Von kossa was performed by staining cultures with 4% silver nitrate solution and exposing cultures to 2400 kJ of ultraviolet light using a UV Stratalinker.

### In vitro Salmon Calcitonin (sCT), Parathyroid Hormone 1-34 (PTH) and Forskolin treatment

WT and *Gnas E1+/−* BMSC or BMM cultures were treated with 10^−7^M of salmon calcitonin (R&D Biosystems), 10^−7^M PTH (Prospec Inc), 10^−5^M of forskolin (Sigma) or vehicle (PBS) controls for 6 hours in serum free αMEM.

### Quantitative RT-PCR

Total RNA was extracted from BMSC and BMM cell cultures, as well as tibia diaphysis tissue samples. For cell culture experiments, adherent cells were rinsed in ice-cold nuclease-free water. Samples were subsequently exposed to 500uL TRIzol (Invitrogen). For tibia harvesting, the tissue samples were freshly dissected, removing the surrounding muscle and connective tissue. After scraping off the periosteum with a 10-blade scalpel, the epiphyses were removed. The remaining tibias were placed within 200uL pipet tip inside of a microcentrifuge tube and underwent centrifugation at 16,000rpm for 60 seconds at 4°C. After removing the marrow, the tibias were placed into RNA Later solution (InVitrogen) and stored at −80°C. For RNA extraction, both tibiae from each mouse were placed into 1 mL of TRIzol (Invitrogen) on ice and homogenized.

To extract RNA, 200uL of Chloroform (Sigma Aldrich) was placed into TRIzol-exposed cell culture or homogenized tissue samples. The clear aqueous layer was isolated and was subsequently run through a Direct-zol RNA Miniprep Kit (Zymo Research). The isolated RNA was treated with DNAse I (New England Bioscience) for 10 minutes at 37°C to remove any remaining genomic DNA, followed by DNAse inactivation by EDTA treatment and heat inactivation at 75°C for 10 minutes. The DNAse-treated RNA samples were then placed through a Monarch RNA Cleanup Kit (NEB) to ensure no carryover of contaminants. 500ng of RNA was utilized for reverse transcription using a high capacity cDNA reverse transcription kit (Applied Biosystems). Quantitative RT-PCR was performed within a 20 μL reaction, consisting of iTaq Universal SYBR Green supermix, 10 μM of forward and reverse primers and 25ng of cDNA. The specific primer sequences utilized are listed below.

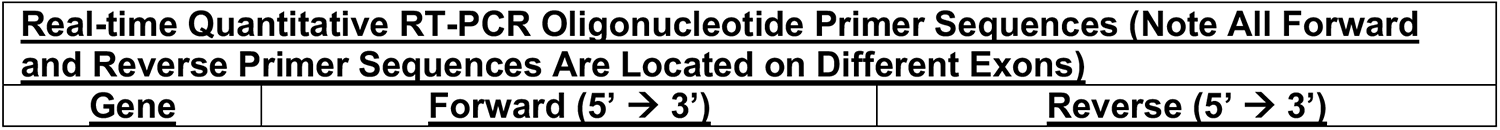
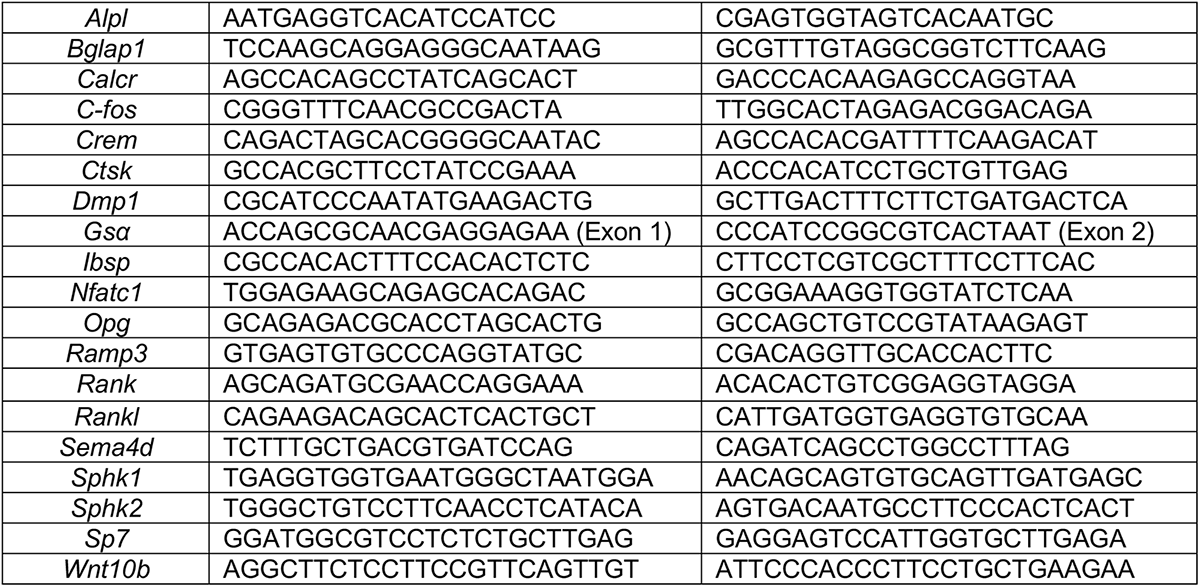

### MCT Analysis

Analysis of femur and lumbar vertebral bone parameters were completed on *Gnas E1+/−* and WT mice using microcomputed tomography (uCT 40; Scanco Medial AG, Bassersdorf, Switzerland). The lumbar vertebrae (L4 through L6) and femurs of WT and *Gnas E1+/−* mice were dissected to remove surrounding connective tissue and fixed in 70% ethanol at 4°C. Cortical bone parameters were measured at the femur mid-diaphysis. Trabecular bone parameters were performed at the distal metaphyseal region of the femur and the centrum of the L5 vertebrae. ΜCT analysis was performed using the following parameters: voxel size of 8μm, 55kV and 145μA.

### Tissue Fixation, Embedding and Sectioning for Bone Histomorphometry

Femur and lumbar vertebrae samples were harvested, immediately dissected to remove surrounding connective tissue and muscle and fixed in 10% Neutral Buffered Formalin (NBF) for 5 days at 4°C. The bones were then briefly transferred to PBS for 2 hours then placed in 30% Sucrose for 48 hours at 4°C. Samples were then embedded into Optimal Cutting Temperature (OCT) and blocks were stored at −80°C until use. Five micrometer (5-μm) undecalcified sections were utilized for histomorphometry analysis._Dynamic histomorphometry was performed by alizarin complexone (AC) and calcein double-labeling, in addition to both Alkaline Phosphatase (AP) and Tartrate-resistant acid phosphatase (TRAP) as previously described.^(48)^ Briefly, WT and *Gnas E1+/−* mice were administered alizarin complexone (AC) by intraperitoneal injection (dose 30mg/kg (Sigma A-3882)) 7 days prior to sacrifice by CO_2_ asphyxiation according to IACUC regulations. These same mice were additionally administered calcein (dose 10mg/kg (Sigma C-0875)) 2 days prior to sacrifice. Tissue specimens of undecalcified distal femur and lumbar vertebrae were imaged using a Ziess Axioscan Z1 high speed automated image acquisition system (Cat#440640-9903-000) and a high resolution camera (AxioCam HRm). Trabecular mineral apposition rate (MAR) analyses, AP and TRAP measurements were calculated using a computer-automated dynamic histomorphometry image analysis software as previously described.^(48)^ Endocortical MAR analysis was performed using the standard histomorphometry analysis software OsteoMetrics (Decatur, GA).

### Serum and Plasma Analyses

Serum and plasma samples were harvested from 12 week old WT and *Gnas E1+/−* mice by cardiac puncture after fasting for 6 hours and stored at −80°C until use. Serum measurements were performed through the use of Immunodiagnostic Systems Mouse Serum Procollagen type 1 N-terminal propeptide (P1NP) (IDS AC-33F1) and Serum Collagen type 1 C-terminal telopeptide (CTX-1) (IDS AC-06F1) ELISA kits. Plasma measurement of calcitonin was performed using an ABClonal Calcitonin Plasma ELISA kit (RK02716). All plasma and serum samples were performed in duplicate and each individual data point generated consisted of the mean value obtained from duplicate samples.

### Statistical Analysis

All statistical analyses were performed using Graphpad Prism Version 9, with *p*-values < 0.05 to be considered as statistically significant. For all analyses observed at once discrete time point, data obtained from WT, *Gnas E1+/−p* and *Gnas E1+/−m* were analyzed by a one-way ANOVA with a post hoc Tukey test for multiple comparisons. For Alizarin Red staining and RT-PCR data comparing changes in mRNA expression at multiple time points or following multiple drug treatments, a two-way ANOVA with a post hoc Tukey test for multiple comparisons was utilized.

## RESULTS

### *Gnas E1+/−m* mice display significantly enhanced cortical and trabecular bone parameters when compared to *Gnas E1+/−p* mice

In order to identify the impact of *Gsα* heterozygous inactivation on cortical and trabecular bone architecture, we utilized our AHO mouse model generated by targeted disruption of exon 1 of *Gnas (Gnas E1+/−)*, leading to global heterozygous inactivation of *Gsα*.^(18)^ We utilized 12-week old mice bred on a pure 129SvEv background, given that this specific background led most closely to recapitulating the human disorder.^(18)^ *Gnas E1+/−m* mice are obese and shorter than their WT counterparts, exhibit hormonal resistance to PTH and TSH, have decreased fertility and develop subcutaneous ossifications.^(18)^ Conversely, *Gnas E1+/−p* mice are not obese and have no hormonal resistance or significant infertility, but they also develop subcutaneous ossifications as in the PPHP human counterpart.^(18,32,33)^ Quantitative RT-PCR of bone-marrow flushed tibia specimens from male and female mice revealed *Gnas E1+/−* mice displayed a significant reduction in *Gsα* mRNA expression when compared to WT (Fig 1A). No significant variations in *Gsα* mRNA expression were observed when comparing *Gnas E1+/−p* to *Gnas E1+/−m* mice (Fig 1A).

**Figure 1:**
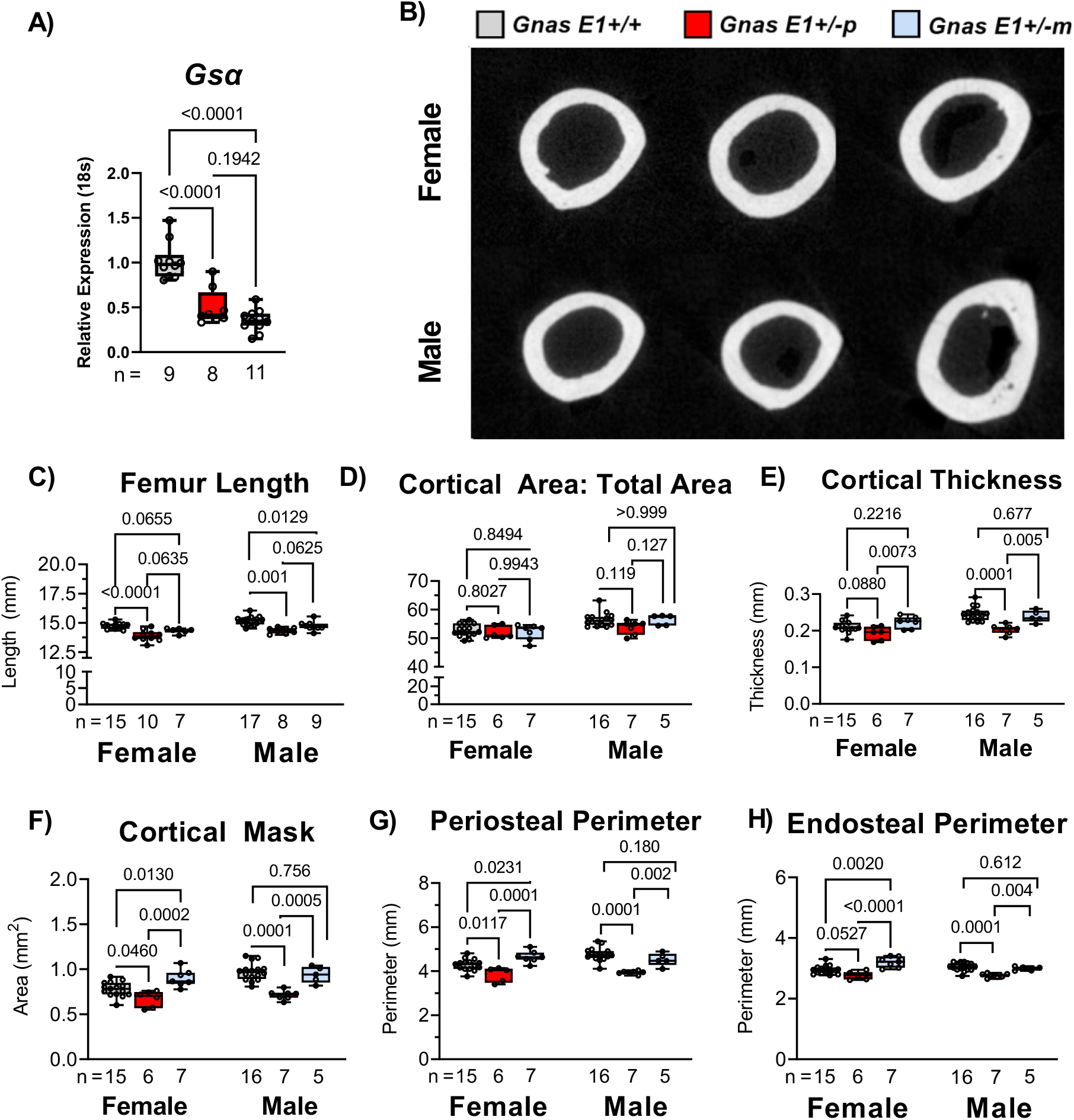
Cortical bone parameters are dependent upon inheritance pattern of *Gsα* mutation. (A) *Gsα* mRNA expression is reduced within flushed tibia samples of *Gnas E1+/−m* and *Gnas E1+/−p* mice compared to WT. (B) Representative cross-sectional images of cortical bone of distal femur across both *WT* and *Gnas E1*+/− animals. (C) Both male and female *Gnas E1+/−p* mice had a significantly reduced femur length compared to *WT*. Male *Gnas E1+/−m* mice also had a significantly reduced femur length compared to WT but no significant variations were observed among female *Gnas E1+/−m* and *WT* mice. (D) Phenotypic differences observed within cortical bone of both *Gnas E1+/−p* and *Gnas E1+/−m* animals are not attributed solely to decreased length, based on the lack of significant differences seen within the ratio of cortical area to total bone area; (E-H) Statistically significant decreases in cortical bone parameters were observed in *Gnas E1+/−p* compared to *Gnas E1+/−m* in both male and female animals. *Gnas E1+/−p* animals demonstrated significant decreases in cortical bone parameters compared to *Gnas E1+/−m* mice, and displayed normal to reduced parameters when compared to *WT*. Conversely, *Gnas E1+/− m* demonstrated normal to increased cortical bone parameters compared to WT. Sample size per genotype per experiment is listed on each bar graph. All statistical tests completed using ANOVA with post-hoc Tukey test for multiple comparisons, and p-values are displayed for each comparison.

The cortical bone architecture of *Gnas E1+/−* and WT mice was analyzed at the mid-diaphysis of the femur by mCT (Fig 1B). All analyses were monitored in both male and female mice, given that previous reports only compared changes in skeletal parameters between male *Gnas E1+/−* and WT mice.^(44)^ Male *Gnas E1+/−* mice (both *Gnas E1+/−p* and *Gnas E1+/−m*) displayed a significant reduction in femur length when compared to WT (Fig 1C); no significant variations were seen in females (*p*=0.06). Despite evidence of reduced femur lengths, *Gnas E1+/− m* and *Gnas E1+/−p* mice displayed no significant variations in the ratio of cortical area to total bone area (CA:TA) when compared to *WT* (Fig 1D), and therefore we were able to make direct comparisons in cortical bone thickness and architecture between each genotype. These analyses revealed significant reductions in cortical bone thickness (Fig 1E), cortical mask (area between periosteal and endosteal surfaces) (Fig 1F) and periosteal- and endocortical-perimeters (Fig 1G-H) in *Gnas E1+/−p* mice when compared to *Gnas E1+/−m* mice and *WT*. These data correlate with earlier findings in our mouse model^(18)^ (including those seen in *Gnas E1+/−* mice bred on a hybrid 129SvEv x CD-1 background) and further confirm that adult *Gnas E1+/−p* mice display reduced cortical bone parameters, whereas *Gnas E1+/−m* mice display normal cortical bone parameters when compared to WT.^(44)^

In conjunction with monitoring changes in cortical bone, we analyzed the trabecular architecture of *Gnas E1+/−* and *WT* mice by MCT at the distal femur (Fig 2A) and lumbar vertebrae (Fig 2B). When comparing the femurs of *Gnas E1+/−p* mice to *WT*, we did not observe a significant trabecular bone phenotype. No significant variations in BV/TV, trabecular number or trabecular spacing were observed in female mice (Fig 2C-F); male *Gnas E1+/−p* mice did display a reduction in trabecular thickness when compared to *WT* but displayed no variations in BV/TV, trabecular number or trabecular spacing. However, the comparison of female *Gnas E1+/−m* mice to *WT* displayed an elevated BV/TV and trabecular number and a reduction in trabecular spacing (Fig 2C-E). No significant variations were observed between male *Gnas E1+/−m* and *WT* mice. When comparing *Gnas E1+/−m* and *Gnas E1+/−p* mice, we observed *Gnas E1+/−m* mice display a significantly elevated increase in trabecular bone thickness (Fig 2D) but displayed no significant variations in BV/TV, trabecular thickness, or trabecular number. MCT analysis of the lumbar vertebrae revealed no significant variations in trabecular bone parameters between *Gnas E1+/− m*, *Gnas E1+/−p* and WT mice (Fig 2G-J). Collectively, these data further highlight the presence of distinctions in both cortical and trabecular bone parameters between adult *Gnas E1+/−m* and *Gnas E1+/−p* mice, particularly within the femur.

**Figure 2:**
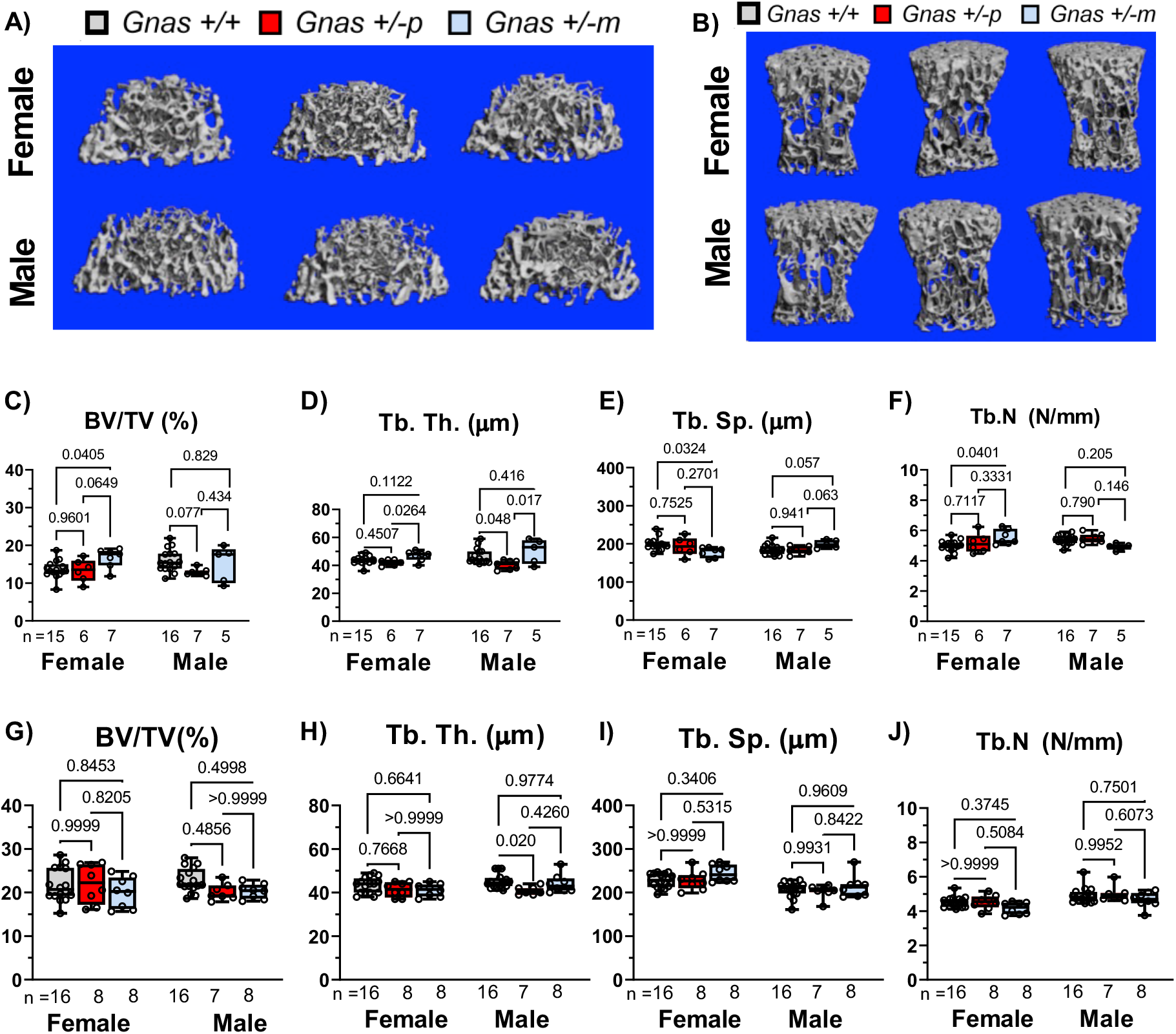
Femur and vertebral trabecular bone parameters are mildly influenced by *Gsα* heterozygous inactivation. Representative μCT images of trabecular bone from the (A) distal femur and (B) L5 vertebrae of WT and *Gnas E1+/−* animals. (C-F) *Gnas E1+/−p* mice, when compared to *WT,* displayed no significant femur trabecular bone phenotype. However, female *Gnas E1+/−m* mice, when compared to *WT* displayed a significantly elevated BV/TV, trabecular number, and a reduction in trabecular spacing (no variations observed between male mice). *Gnas E1+/−m* mice, when compared to *Gnas E1+/− p*, displayed a significant increase in trabecular thickness. (G-J) No significant differences were observed between *WT* and *Gnas E1+/−* mice with respect to vertebral trabecular bone parameters. Sample size per genotype per experiment is listed on each bar graph. All statistical tests completed using ANOVA with post-hoc Tukey test for multiple comparisons, and p-values are displayed for each comparison.

### Elevated cortical and trabecular bone parameters in *Gnas E1+/−m* mice are associated with increased osteoblastic activity *in vivo*

We next evaluated whether the observed differences in cortical and trabecular bone between *Gnas E1+/−m* and *Gnas E1+/−p* mice could be attributed to variations in osteoblast number or underlying osteoblast activity. To address these questions, we utilized a computer-automated fluorescent-based multiplex histomorphometry technique to label actively-mineralizing osteoblasts along the bone surface of the distal femur and lumbar vertebrae using alkaline phosphatase staining in conjunction with calcein and alizarin complexone mineral labeling (Fig 3A).^(48)^ In addition to histomorphometry, we compared overall bone forming activity *in vivo* by serum P1NP measurements and gene expression analyses of osteoblast-related transcripts osteocalcin (*Bglap1*), integrin-binding sialoprotein (*Ibsp*) and dentin matrix acidic phosphoprotein 1 (*Dmp1*) by RT-PCR.

**Figure 3:**
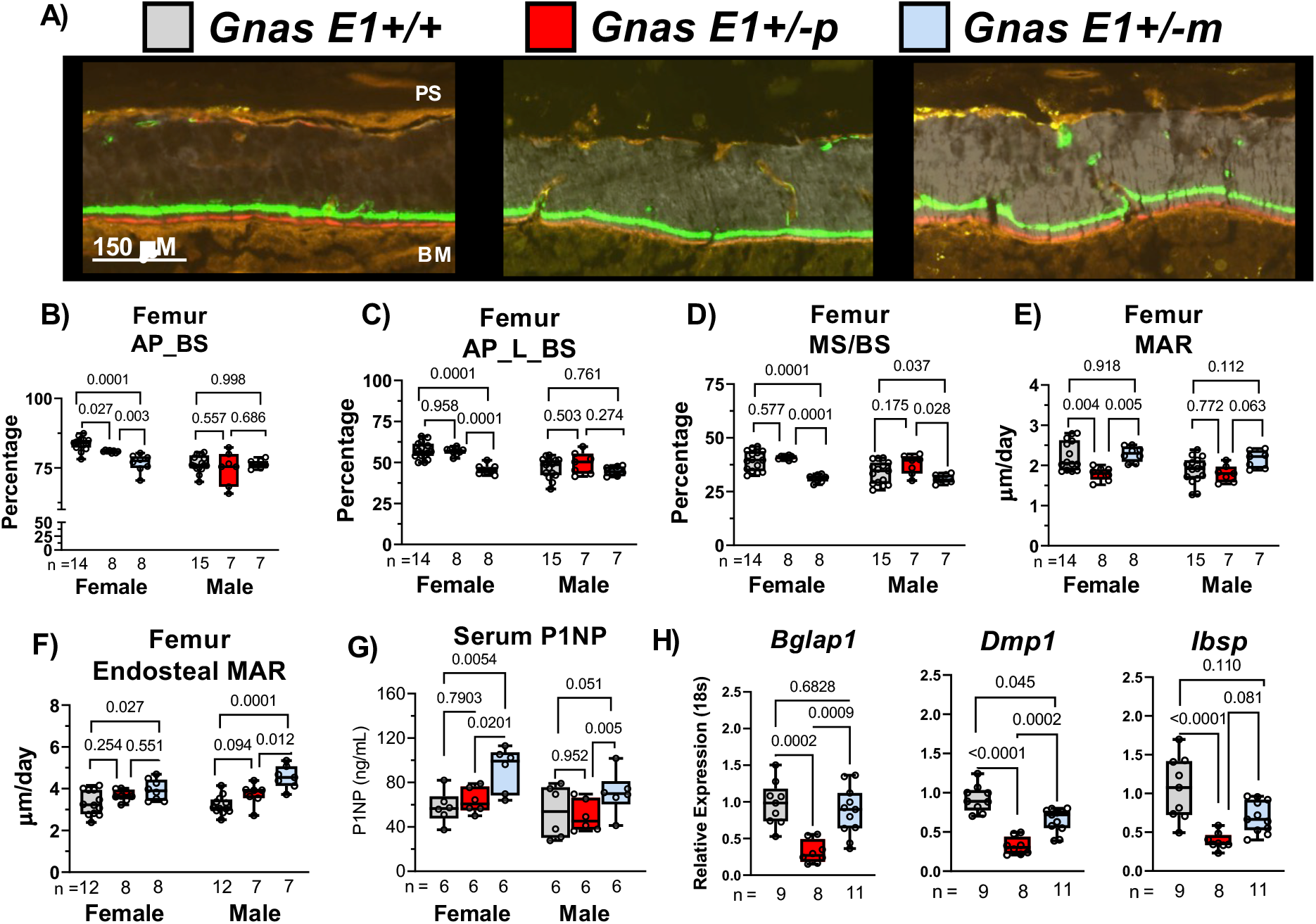
Enhanced cortical and trabecular bone parameters in *Gnas E1+/−m* mice are directly associated with elevated osteoblast activity despite a reduction in osteoblast number. (A) Representative images of undecalcified femur cortical bone sections containing calcein (green) and alizarin complexone (red) labels for dynamic histomorphometry. (B) Significant reductions in total alkaline phosphatase activity on the bone surface (AP_BS) were observed within female *Gnas E1+/− p* and *Gnas E1+/−m* femur sections compared to WT. No significant differences were observed in male specimens across all genotypes. (C) *Gnas E1+/−m* mice displayed a reduction in the total number of actively mineralizing osteoblasts (AP_L_BS) when compared to WT and *Gnas E1+/−p* mice. (D) Dynamic histomorphometry on the femur trabecular surface revealed *Gnas E1+/−m* mice displayed a significant reduction in MS/BS within the femur when compared to both *WT* and *Gnas E1+/−p* mice (E) Female *Gnas E1+/−p* mice had a significantly reduced mineral apposition rate compared to both *Gnas E1+/−m* and WT. MAR among *Gnas E1+/−m* was comparable to WT. We observed no significant variations within male mice. (F) *Gnas E1+/−m* mice displayed a significantly increased MAR on the femoral endosteal surface compare to WT mice, as well as when compared to *Gnas E1+/−p* mice for males. (G) Serum P1NP levels were significantly elevated within *Gnas E1+/−m* mice compared to *Gnas E1+/−p* and WT samples. No significant variations were observed between *Gnas E1+/−p* and WT samples. (H) *Bglap1, Dmp1* and *Ibsp* mRNA expression in flushed tibia specimens was significantly reduced in *Gnas E1+/−p* male and female mice compared to WT. *Gnas E1+/−m* samples displayed no significant reduction in *Bglap, Ibsp* expression when compared to WT but displayed a reduction in *Dmp1* expression. *Gnas E1+/−m* samples, when compared to *Gnas E1+/−p,* displayed a significantly elevated *Bglap1* and *Dmp1* expression, but displayed no significant variations in *Ibsp* expression. Sample size per genotype per experiment is listed on each bar graph. All statistical tests completed using ANOVA with post-hoc Tukey test for multiple comparisons, and p-values are displayed for each comparison.

While histomorphometric analyses were performed using both male and female mice, we primarily observed significant variations in overall bone remodeling within female *Gnas E1+/−* and WT mice. This dimorphism is aligned with previous studies that have demonstrated within the skeleton of over 220 mouse lines, female mice display higher trabecular bone turnover and therefore may be more susceptible to phenotypic perturbations when compared to male mice.^(49)^ In particular, we observed both female *Gnas E1+/−m* and *Gnas E1+/−p* mice displayed a statistically significant reduction in the presence of osteoblasts along the bone surface (AP_BS) when compared to WT at both the distal femur (Fig 3B) and lumbar vertebrae (Supplemental Figure 1A). Despite female *Gnas E1+/−m* mice displaying enhanced cortical and trabecular bone architecture, these mice displayed a reduction in the number of actively mineralizing osteoblasts (measured by both AP+ populations superimposed over a mineralization line (AP_L_BS) and MS/BS) within the distal femur (Fig 3C,D) and lumbar vertebrae (Supplemental Figure 1B,C) when compared to both *Gnas E1+/−p* and WT mice.

We utilized calcein and alizarin complexone double labeling to quantify the mineral apposition rate (MAR) along three separate bone surfaces in *Gnas E1+/−* and WT mice: the trabecular surfaces of the distal femur, the endocortical surface of the femur, and the trabecular surface of the lumbar vertebrae. Along the femur trabecular surface, female *Gnas E1+/−p* mice displayed a significant reduction in MAR when compared to both *Gnas E1+/−m* and WT (Fig 3E). Alternatively, *Gnas E1+/−m* mice displayed MAR values that were comparable to WT despite displaying an overall reduction in the number of active osteoblasts on the bone surface. No significant variations in trabecular MAR were observed in male mice (Fig 3E). Subsequent analysis of MAR along both the femur endocortical surface (Fig 3A, 3F) and the lumbar vertebrae trabecular surface (Supplemental Figure 1D) revealed that *Gnas E1+/−m* mice displayed a significantly elevated MAR compared to WT, whereas no significant differences were observed between *Gnas E1+/−p* and WT mice at either of these two locations. Complementary to our histomorphometry findings, female *Gnas E1+/−m* mice displayed significantly elevated serum P1NP measurements when compared to both *Gnas E1+/− p* and WT (Fig 3G). Male *Gnas E1+/− m* mice also displayed elevated P1NP levels when compared to *Gnas E1+/−p* mice, but were not significantly higher than WT (*p*=0.*051*). No significant differences were observed between *Gnas E1+/−p* and WT mice in both males and females. Quantitative RT-PCR analysis of flushed tibia specimens from male and female mice displayed *Gnas E1+/−p* mice having a significant reduction in *Bglap1 (p*<0.001)*, Dmp1 (p<0.001*) and *Ibsp* (*p<0.001*) (Fig 3H) mRNA expression when compared to WT. These data complement our histologic findings in the femur and lumbar vertebrae and further suggest evidence of reduced osteoblast activity within *Gnas E1+/−p* mice compared to WT. However, *Gnas E1+/−m* mice displayed no significant variations in *Bglap1* (*p=*0.*683*) and *Ibsp (p=0.110*) mRNA expression when compared to WT but did have a reduction in *Dmp1* expression (*p=0.045*). In addition, *Gnas E1+/−m* mice displayed a significant increase in *Bglap1* (*p=0.0009*) and *Dmp1* (*p=0.0002*) expression when compared to *Gnas E1+/−p* but displayed no significant variation in *Ibsp (p=0.081*) expression. These data suggest that despite an overall reduction in the total number of active osteoblasts within the femoral and vertebral bone surface of *Gnas E1+/−* mice, the enhanced cortical and trabecular bone parameters specifically observed within *Gnas E1+/−m* mice can be attributed to an increase in the total matrix production per osteoblast *in vivo* when compared to *Gnas E1+/−p* mice.

Given these observed distinctions in osteoblast activity *in vivo*, we next isolated primary bone marrow stromal cells (BMSCs) from both male and female mice to identify whether there were any significant variations in the transcriptional activity or osteogenic differentiation within the osteoprogenitor populations of *Gnas E1+/−m* and *Gnas E1+/−p* mice (Fig 4). When cultured in basal media (αMEM with 10% FBS), no significant variations were observed between the osteoprogenitor populations of *Gnas E1+/−* and WT cultures, as measured by the number of AP+ colony forming units (Fig 4A-B). When supplemented with osteogenic inducing media for 7 and 14 days, no significant variations in osteogenic differentiation or mineralization were observed between *Gnas E1+/−* and *WT* cultures as measured by alizarin red or von Kossa staining (Fig 4A,C). Gene expression analyses of *Gsα* and osteogenic-related genes *Alpl* (Alkaline phosphatase), *Sp7* (Osterix) and *Bglap1* were also performed within cultures in basal media (Day 0) and following 7 and 14 days of osteogenic differentiation (Fig 4D-G). At each timepoint, both *Gnas E1+/−m* and *Gnas E1+/−p* cultures displayed significant reductions in *Gsα* mRNA expression (Fig 4D) when compared to *WT* cultures. When cultured in basal media, no significant variations in *Alpl*, *Sp7* or *Bglap1* were observed between *Gnas E1+/−* and *WT* cultures (Fig 4E-G). However, *Gnas E1+/−m* cultures after 7 days of osteogenic differentiation displayed a significant upregulation in *Alpl* (Fig 4E) and *Sp7* (Fig 4F) mRNA expression when compared to WT and *Gnas E1+/−p*. Additionally, after 14 days of osteogenic differentiation, *Gnas E1+/−m* BMSCs displayed an increase in *Alpl* (Fig 4E) and *Bglap1* (Fig 4G) mRNA expression compared to WT and *Gnas E1+/−p* cultures. While *Gnas E1+/−m* BMSCs display elevated osteogenic transcriptional activity *in vitro*, the lack of a mineralization or differentiation phenotype *in vitro* suggests the observed changes in osteoblast activity within *Gnas E1+/−m* mice *in vivo* may not be solely driven by cell autonomous changes in osteogenic differentiation capacity. Rather, these observed *in vivo* findings could alternatively be driven by local changes within the bone microenvironment.

**Figure 4:**
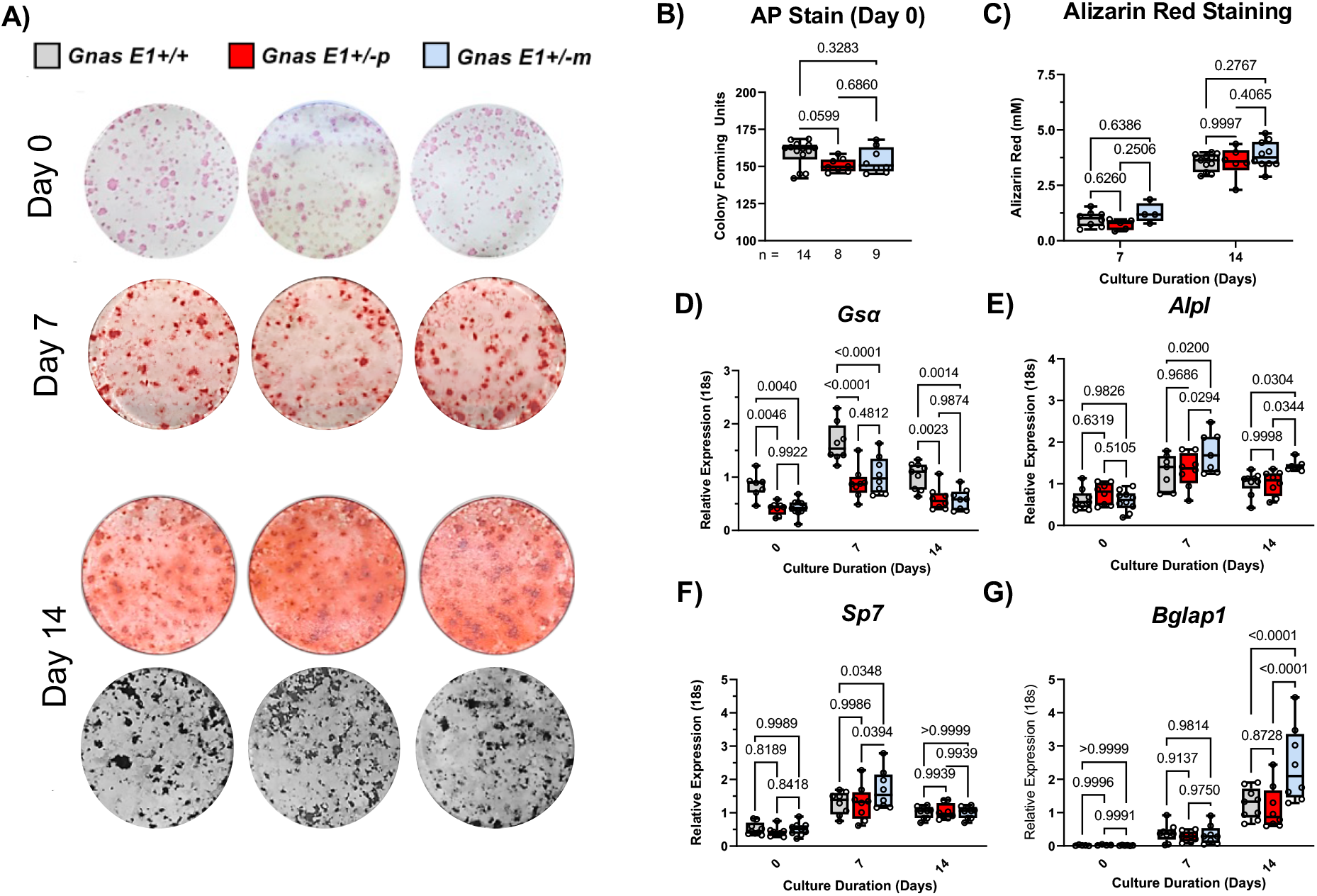
Inheritance pattern of *Gsα* mutation does not differentially influence osteoprogenitor differentiation *in vitro*. (A) Representative entire-well images of: Alkaline Phosphatase positive (AP+) colony formation units within primary BMSCs at day 0 from male and female mice; Alizarin Red staining to detect calcium deposition on culture days 7 and 14; and Von kossa staining to detect phosphate on culture day 14. (B) Both *Gnas E1+/−m* and *Gnas E1+/−p* demonstrated no significant differences within the osteoprogenitor populations compared to WT BMSCs as measured by alkaline phosphatase colony forming unit assays. (C) *Gnas E1+/−* and WT BMSCs following 7 and 14 days of osteogenic differentiation displayed no significant variations in alizarin red staining absorbance. (D) Both *Gnas E1+/−m* and *Gnas E1+/−p* BMSCs displayed significant reductions in *Gsα* expression at each time point when compared to *WT.* Prior to osteogenic differentiation, no significant differences were observed in *Alpl, Sp7* and *Bglap1* mRNA expression among *Gnas E1+/−* and WT BMSCs. However, *Gnas E1+/−m* mice displayed elevated mRNA expression of *Alpl, Sp7* after 7 days of osteogenic differentiation and *Alpl* and *Bglap1* after 14 days of osteogenic differentiation when compared to both WT and *Gnas E1+/−p*. Sample size per genotype per experiment is listed on each bar graph. For AP+ colony assay, statistical tests were completed using a one-way ANOVA with post-hoc Tukey test for multiple comparisons. For alizarin red staining and RT-PCR analyses at multiple timepoints, statistical tests were completed using a two-way ANOVA with post-hoc Tukey test for multiple comparisons. P-values are displayed for each comparison.

### *Gsα* heterozygous inactivation differentially affects osteoclast-secreted anabolic coupling factors *in vitro*

Because *Gnas E1+/−m* mice displayed elevated bone forming activity *in vivo* but did not display variations in osteogenic differentiation or mineralization capacity *in vitro,* we next investigated whether these distinctions between *Gnas E1+/−m* and *Gnas E1+/−p* mice could be driven by modifications within the bone microenvironment. For this approach, we first assessed whether *Gsα* heterozygous inactivation differentially influences osteoclast differentiation and structural morphology. *In vitro* analysis of primary bone marrow macrophage (BMM) cultures from male and female mice revealed both *Gnas E1+/−m* and *Gnas E1+/−p* cultures had elevated rates of osteoclastogenesis when compared to WT as measured by chromogenic tartrate-resistant acid phosphatase (TRAP) staining (Fig 5A-B). Fluorescent microscopy using phalloidin revealed *Gnas E1+/−m* and *Gnas E1+/−p* osteoclasts displayed appropriate actin ring morphology and were significantly larger in overall surface area when compared to WT (Fig 5A-B). Quantitative RT-PCR revealed *Gnas E1+/−m* and *Gnas E1+/−p* BMM cultures displayed a significant upregulation of *Rank* (Fig 5C) and *Calcr* (Fig 7A) mRNA expression when compared to WT, but displayed no significant variations in *Nfatc1* or *Ctsk* mRNA expression (Fig 5C). Collectively, these data correlate with previous reports suggesting *Gsα* serves as a negative regulator of osteoclastogenesis and that parental inheritance of the *Gsα* mutation does not selectively influence osteoclastogenesis *in vitro*.^(44,50)^

**Figure 5:**
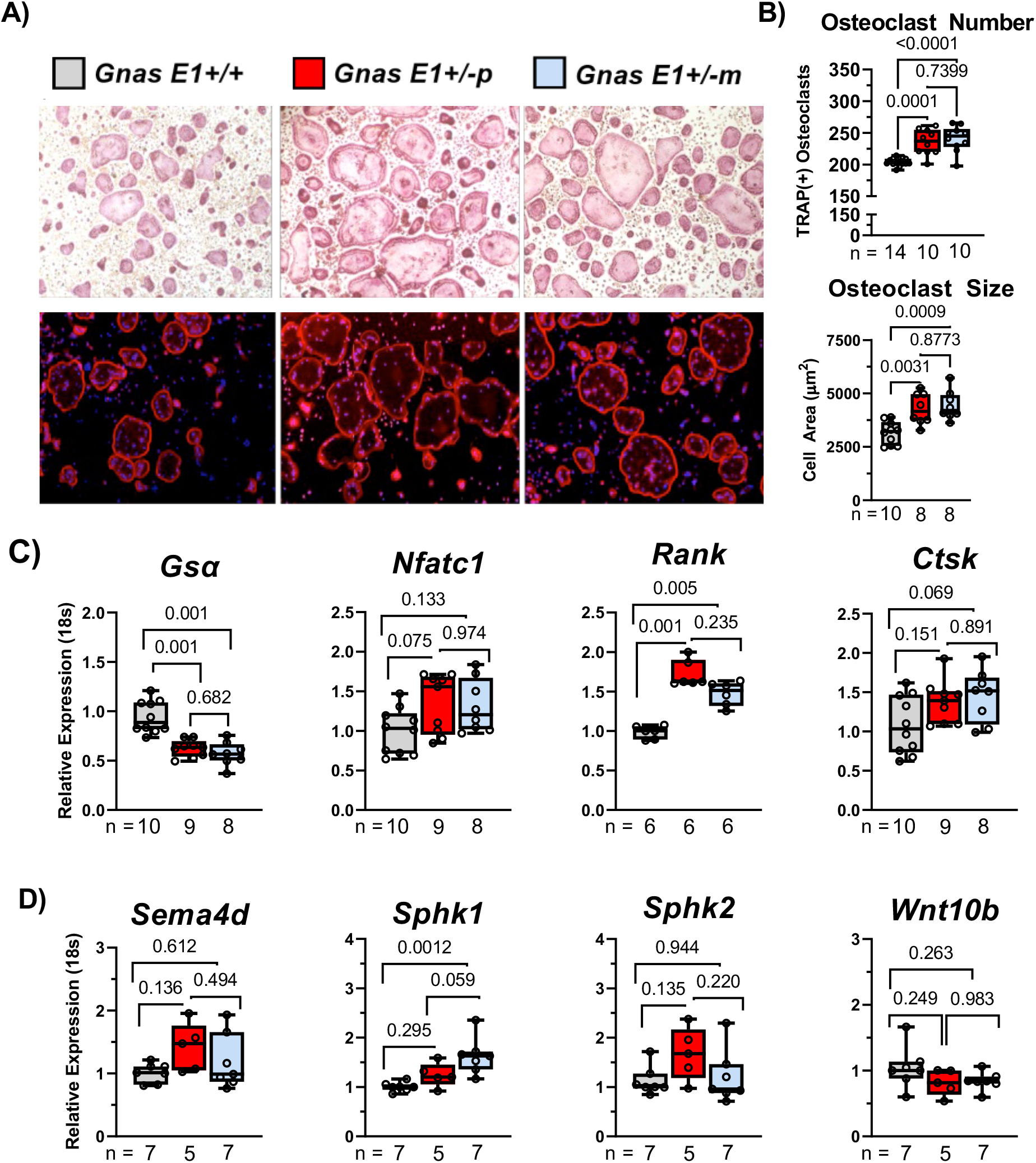
*Gnas E1+/−m* osteoclasts display elevated *Sphk1* expression *in vitro*. (A) Representative images of chromogenic TRAP staining (top) and phalloidin-DAPI fluorescence microscopy staining of primary BMM cultures (bottom) from male and female mice *in vitro*. (B) *Gnas E1+/−m* and *Gnas E1+/−p* demonstrated increased rates of osteoclastogenesis and osteoclast surface area compared to WT as indicated by increased osteoclast number following 5 days of differentiation. Osteoclasts were defined as TRAP(+) cells with ≥3 nuclei. No significant differences were observed in osteoclast number between *Gnas +/−p* and *Gnas +/−m* cultures, respectfully; (C) *Gsα* mRNA expression was significantly reduced in both *Gnas E1+/−m* and *Gnas E1+/−p* cultures when compared to WT. No significant variations were observed in mRNA expression of *Nfatc1* or *Ctsk* among *Gnas E1+/−* cultures and WT at day 5 of osteoclast differentiation. However, both *Gnas E1+/−p* and *Gnas E1+/−m* cultures demonstrated a significant increase in *Rank* expression compared to WT. (D) *Gnas E1+/−m* cultures display significantly elevated *Sphk1* mRNA expression when compared to WT cultures. Sample size per genotype per experiment is listed on each bar graph. All statistical tests completed using ANOVA with post-hoc Tukey test for multiple comparisons, and p-values are displayed for each comparison.

In addition to evaluating osteoclastogenesis *in vitro*, we wanted to assess whether the observed distinctions in bone formation between *Gnas E1+/−m* and *Gnas E1+/−p* mice *in vivo* could be attributed to modifications in osteoclast secreted bone coupling factors. We examined four growth factors that are released by osteoclasts and have been shown to influence bone formation: *Semaphorin 4d* (*Sema4d*), *Sphingosine Kinase 1 (Sphk1), Sphingosine Kinase 2 (Sphk2)* and *Wnt10b* (Fig 5D).^(51–56)^ *Gnas E1+/−p* BMM cultures displayed no significant variations in transcriptional activity when compared to WT. However, *Gnas E1+/−m* BMM cultures displayed significantly enhanced *Sphk1* transcriptional activity when compared to *WT* (*p=0.001*), but were not statistically significant when compared to *Gnas E1+/−p* cultures (*p=0.059*) (Fig 5D). *Sphk1* and Sphingosine 1-phosphate (S1P) signaling within the osteoclast has been well recognized to promote osteoblast function *in vivo*.^(51–56)^ Therefore, these data underscore that the distinctions in bone forming activity between *Gnas E1+/−m* and *Gnas E1+/−p* mice could be attributed to variations in osteoblast-osteoclast coupling interactions within the bone microenvironment secondary to S1P-related signaling.

### Reduced cortical and trabecular bone parameters within *Gnas E1+/−p* are attributed to enhanced osteoclast activity *in vivo*

In conjunction with assessing bone formation, we compared overall bone resorption between *Gnas E1+/−* and WT mice *in vivo* by histomorphometry at the distal femur and vertebrae, serum CTX-1 measurement and RT-PCR to measure *Rankl* and *Opg* mRNA expression within flushed tibia samples. Histomorphometric analysis to assess osteoclast activity was performed by incubating sections with Elf97 substrate (Life Tech, E6589), which generates a fluorescent signal when cleaved by TRAP and has been reproducibly shown to stain osteoclasts *in vivo* (Fig 6A).^(48,57)^

**Figure 6:**
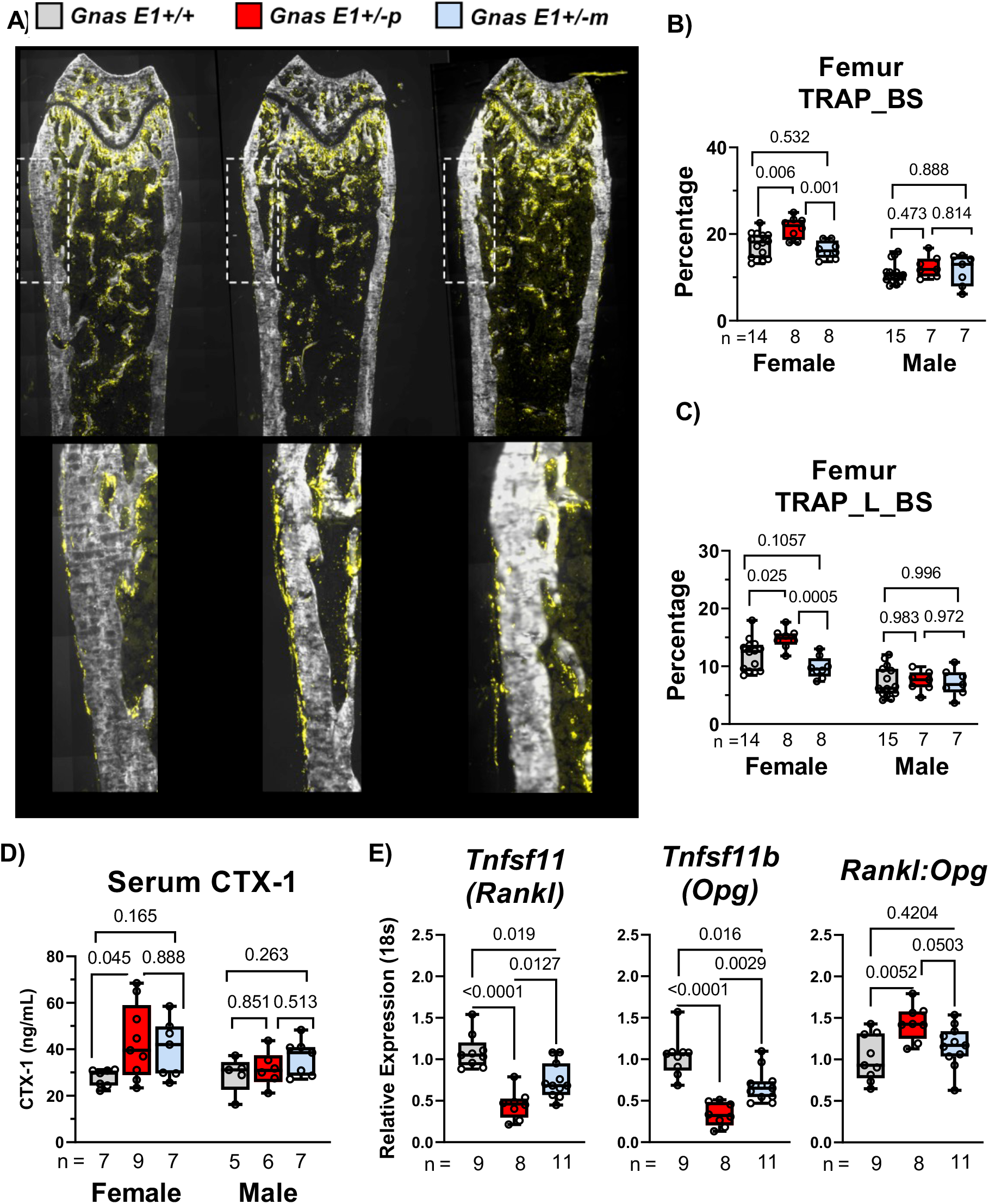
*Gsα* heterozygous inactivation differentially affects bone resorption in a parental inheritance-specific pattern *in vivo*. (A) Representative images of undecalcified distal femur sections demonstrating TRAP enzymatic activity on the bone surface (TRAP_BS) following incubation with Elf97 fluorescent substrate. Female *Gnas E1+/−p* mice demonstrated a significant increase in (B) total TRAP activity on the bone surface (TRAP_BS) and (C) active bone remodeling sites (TRAP_L_BS) within the femur when compared to WT and *Gnas E1+/−m* mice. No significant variations were observed between female *Gnas E1+/−m* and WT mice. No significant phenotype was observed within male mice. (D) Female *Gnas E1+/−p* mice displayed significantly elevated fasting serum CTX-1 measurements when compared to WT. (E) RT-PCR analysis of flushed tibia diaphysis from male and female mice identified both *Gnas E1+/− p* and *Gnas E1+/−m* mice had reduced mRNA expression of *Tnfsf11* (Rankl) and *Tnfsf11b* (Opg) when compared to WT. *Gnas E1+/−m* mice displayed significantly increased *Rankl* and *Opg* mRNA expression when compared to *Gnas E1+/−p.* However, *Gnas E1+/−p* mice displayed a significantly elevated *Rankl:Opg* ratio when compared to WT but were not statistically significant when compared to *Gnas E1+/−m* mice. Sample size per genotype per experiment is listed on each bar graph. All statistical tests completed using ANOVA with post-hoc Tukey test for multiple comparisons, and p-values are displayed for each comparison.

Histologic analyses at the distal femur revealed that *Gnas E1+/−p* mice display elevated TRAP activity on the periosteal surface when compared to both *WT* and *Gnas E1+/−m* mice (Fig 6A). In addition, female *Gnas E1+/−p* mice displayed elevated total TRAP activity (TRAP_BS) (Fig 6B) and active remodeling surfaces (defined as TRAP+ signals over a mineralizing surface, or TRAP_L_BS) (Fig 6C) when compared to both *Gnas E1+/−m* and WT. No significant variations were observed between female *Gnas E1+/−m* and WT samples. Within the lumbar vertebrae, *Gnas E1+/−p* mice displayed an elevated TRAP_BS when compared to WT and had an elevated TRAP_L_BS when compared to both *WT* and *Gnas E1+/−m* mice (Supplemental Figure 1E,F). Complementary to these histomorphometry findings, female *Gnas E1+/−p* mice displayed significantly elevated serum CTX-1 compared to WT (Fig 6D), whereas no significant changes were observed in *Gnas E1+/−m* serum levels. Additionally, no significant variations were observed between *Gnas E1+/−p* and *Gnas E1+/−m* measurements. These data correlate with previous reports in the literature that demonstrate enhanced osteoclast activity specifically among *Gnas E1+/−p* mice.^(44)^

Quantitative RT-PCR from tibia specimens isolated from both male and female mice revealed that *Gnas E1+/−p* and *Gnas E1+/−m* mice displayed a significant reduction in both *Rankl* and *Opg* (Fig 6E) mRNA expression when compared to WT. Despite both *Rankl* and *Opg* being downregulated in *Gnas E1+/−* mice, *Gnas E1+/−p* mice observed a more significant reduction in *Opg* expression when compared to *Rankl* reduction which directly correlates with previous tibiae and calvaria mRNA data in the literature from *Gsα* deficient animals.^(36,37)^ Consequently, this resulted in *Gnas E1+/−p* mice displaying an elevated *Rankl:Opg* ratio (Fig 6E) when compared to WT. *Gnas E1+/−m* mice exhibited a more modest reduction in both *Rankl* and *Opg* expression when compared to WT, with no significant change in the *Rankl:Opg* ratio. Collectively, despite *Gnas E1+/−m* and *Gnas E1+/−p* mice displaying equally elevated rates of osteoclastogenesis *in vitro,* only *Gnas E1+/−p* mice demonstrate elevated bone resorption activity *in vivo*. These data demonstrate that the reduced cortical and trabecular bone parameters within *Gnas E1+/−p* mice are driven by a combination elevated resorption activity specifically within *Gnas E1+/−p* mice, in conjunction with their impaired bone formation activity.

### Inheritance -specific variations in bioactivity of Gsα-coupled receptors are observed in calcitonin signaling within the osteoclast lineage among *Gnas E1+/−* mice *in vitro*

In light of these distinctions in bone remodeling among *Gnas E1+/−m* and *Gnas E1+/−p* mice, we next sought to identify whether these differences could be driven by changes in bioactivity of Gsα-coupled receptors within specific cell-types in the bone microenvironment. Although multiple hormones have been identified to influence bone homeostasis through Gsα-mediated signaling, we were particularly interested in comparing the responsiveness of osteoclasts to the hormone calcitonin, as well as observing the responsiveness of osteoblasts to parathyroid hormone (PTH). It is well-described that patients with PHP1A can manifest resistance to calcitonin.^(45,46)^ In primary BMM cultures, we observed a significant upregulation in Calcitonin receptor (*Calcr*) mRNA expression in both *Gnas E1+/−m* and *Gnas E1+/−*p cells compared to WT (Fig 7A). Given that *Calcr* is an often-utilized transcriptional marker of mature osteoclasts *in vitro*, this finding was anticipated based upon the increased rates of osteoclastogenesis in both *Gnas E1+/−m* and *Gnas E1+/−p* cultures. However, *Gnas E1+/−m* primary cultures also displayed a consistent upregulation of *Calcr* expression when compared to *Gnas E1+/−p* cultures despite having similar numbers of osteoclasts *in vitro*.

**Figure 7:**
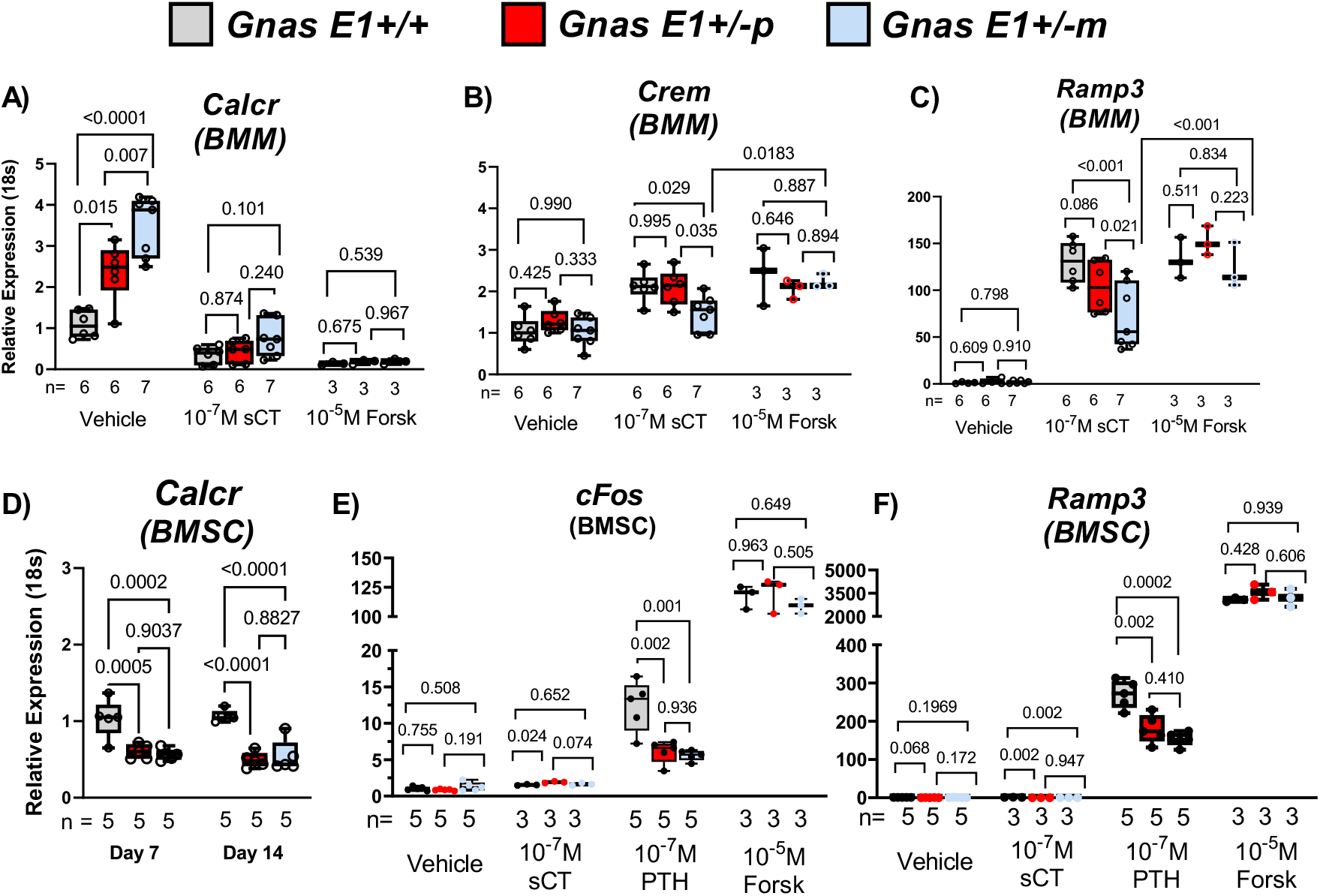
*Gnas E1+/−m* osteoclasts display impaired calcitonin receptor bioactivity *in vitro.* (A) *Gnas E1+/−m* and *Gnas E1+/−p* osteoclasts displayed significant elevations in *Calcr* mRNA expression when compared to WT; however, *Calcr* expression within *Gnas E1+/−m* cultures was significantly greater than *Gnas E1+/−p* cultures. Treatment of *Gnas E1+/−* and WT osteoclasts with 10^−7^ M salmon CT or 10^−5^ M forskolin resulted in significant reductions in *Calcr* expression. (B-C) RT-PCR analysis of *Crem* and *Ramp3* mRNA expression of *Gnas E1+/−m, Gnas E1+/−p* and WT cultures following exposure to salmon CT or vehicle controls for 6 hours. sCT treated *Gnas E1+/−m* cultures displayed a significant reduction in *Crem* and *Ramp3* mRNA expression compared to *Gnas E1+/−p* and WT sCT treated cultures. No significant variations in *Crem* or *Ramp3* expression were observed between *Gnas E1+/−p* and WT sCT treated cultures. (D) *Gnas E1+/−m* and *Gnas E1+/−p* BMSCs following 7 and 14 days of osteogenic differentiation demonstrate a significant reduction in *Calcr* mRNA expression when compare to WT. (E-F) RT-PCR analysis of *c-Fos* and *Ramp3* mRNA expression of *Gnas E1+/−m, Gnas E1+/−p* and WT BMSCs following exposure to salmon CT, PTH, forskolin or vehicle controls for 6 hours. BMSCs overall displayed no significant response to sCT treatment. However, *Gnas E1+/−m* and *Gnas E1+/−p* BMSCs displayed a reduction in *c-Fos* and *Ramp3* when compared to WT BMSCs following PTH treatment. No significant variations in *c-Fos* or *Ramp3* expression were observed between PTH treated *Gnas E1+/−m* and *Gnas E1+/−p* BMSCs. Sample size per genotype per experiment is listed on each bar graph. All statistical tests completed using two-way ANOVA with post-hoc Tukey test for multiple comparisons, and p-values are displayed for each comparison.

Within the osteoclast, the transcriptional activity of *Calcr* is maintained through a negative autoregulatory feedback loop based upon calcitonin receptor activation.^(58,59)^ We therefore wanted to address whether the observed upregulation in *Calcr* expression specifically within *Gnas E1+/− m* cultures was attributed to impaired calcitonin receptor Gsα-mediated signaling. *Gnas E1+/−m, Gnas E1+/−p*, and WT BMMs were treated with 10^−7^M salmon calcitonin (R&D biosystems) to activate Gsα signaling, with 10^−5^M forskolin (Sigma) to activate adenylyl cyclase (AC) downstream of Gsα, or with vehicle controls (PBS) for 6 hours. We monitored calcitonin receptor bioactivity by quantitative RT-PCR analysis of *Calcr*, as well as *Crem* and *Ramp3*. These two additional genes were selected because their transcriptional activity has been shown to be directly influenced by signaling through Gsα-coupled receptors and have been previously validated to monitor the responsiveness of osteoclasts to calcitonin *in vitro*.^(52,60)^ Exposure of *WT, Gnas E1+/−p*, and *Gnas E1+/−m* cultures to sCT and forskolin resulted in a reduction of *Calcr* mRNA expression when compared to vehicle-treated cultures (Fig 7A). We did not observe any statistically significant variations in *Calcr* expression among sCT treated cultures*. Gnas E1+/−m* osteoclast cultures treated with sCT displayed a significant reduction of both *Crem* (Fig 7B) and *Ramp3* (Fig 7C) mRNA expression when compared to both WT and *Gnas E1+/−p* sCT treated cultures, as well as forskolin-treated *Gnas E1+/−m* cultures. Both *Crem* and *Ramp3* mRNA expression levels in *Gnas E1+/−p* cultures following sCT treatment were comparable to WT. In addition, treatment of cultures with forskolin displayed no significant variations in *Crem* or *Ramp3* mRNA expression between *Gnas E1+/−* or WT treated cultures. Therefore, these data suggest evidence of partial calcitonin resistance within *Gnas E1+/−m* osteoclasts *in vitro* and imply that this reduced activity is driven specifically by impaired calcitonin receptor Gsα-mediated signaling. Although we did not observe any variations in *Gsα* transcription between *Gnas E1+/−p* and *Gnas E1+/−m* BMM cultures *in vitro* (Fig 5C), these statistical changes in calcitonin-receptor bioactivity may be reflective of a preferential targeting of osteoclasts within this mixed culture system and further investigations are necessary.

### Bialleleic expression of *Gsα* within the osteoblast lineage in *Gnas E1+/−* mice *in vitro*

Given these distinctions in *Calcr* mRNA expression and receptor bioactivity in osteoclast cultures, we subsequently evaluated *Calcr* expression within BMSCs following 7 and 14 days of osteogenic differentiation (Fig 7D), as well as in flushed tibia samples (Supplemental Fig 2A). We observed that BMSCs from *Gnas E1+/−m* and *Gnas E1+/−p* mice displayed significant reductions in *Calcr* mRNA expression when compared to *WT* after 7 and 14 days of osteogenic differentiation (Fig 7D). Similarly, flushed tibia samples from *Gnas E1+/−m* and *Gnas E1+/−p* mice displayed significant reductions in *Calcr* mRNA expression when compared to *WT* (Supplemental Fig 2A). Although these data would suggest similarities in overall Gsα-receptor mediated signaling within the osteoblast, we wanted to more closely confirm this finding *in vitro* by exposing BMSCs isolated from *Gnas E1+/−* and *WT* mice that were treated for 14 days in osteogenic differentiation media to 10^−7^M salmon calcitonin (R&D biosystems), 10^−7^ PTH 1-34 (Prospec), 10^−5^M forskolin (Sigma), or vehicle controls (PBS) for 6 hours. We assessed the bioactivity of Gsα-coupled receptors within BMSCs by monitoring the mRNA expression of *c-Fos, Ramp3* and *Crem* which have been previously validated in the literature to identify Gsα-mediated signaling within BMSCs particularly in response to treatment with PTH.^(61)^ Treatment of both *Gnas E1+/−* and *WT* BMSCs with salmon calcitonin resulted in no significant changes in *c-Fos* (Fig 7E)*, Ramp3* (Fig 7F) or *Crem* (Supplemental figure 2B) expression when compared to vehicle treated cultures. This finding was expected based upon previous reports documenting an inability of osteoblasts to respond to calcitonin despite their mRNA expression of *Calcr*.^(52,62)^ Treatment of *Gnas E1+/−* and *WT* BMSCs with forskolin resulted in a robust increase in *c-Fos, Ramp3* and *Crem* expression when compared to vehicle treated controls; additionally, no significant variations in mRNA expression were observed when comparing forskolin treated *Gnas E1+/−m, Gnas E1+/−p*, and *WT* cultures. Treatment of *Gnas E1+/−* and *WT* BMSCs with PTH 1-34 also caused an upregulation in *Ramp3* and *cFos* mRNA expression when compared to vehicle treated controls (Fig 7E, 7F). Significant changes in *Crem* expression compared to vehicle controls were only observed in *WT* cultures (Sup Fig 2B). When comparing PTH treated BMSCs, both *Gnas E1+/−m* and *Gnas E1+/−p* BMSCs displayed a significant reduction in both *c-Fos* and *Ramp3* mRNA expression when compared to *WT.* No significant variations in *Ramp3, cFos* or *Crem* were observed when comparing *Gnas E1+/−p* to *Gnas E1+/−m* PTH-treated cultures. Collectively, the combination of similar reductions in *Gsα* and *Calcr* mRNA expression between *Gnas E1+/−m* and *Gnas E1+/−p* BMSCs when compared to WT, in addition to both *Gnas E1+/−m* and *Gnas E1+/−p* BMSCs displaying impaired Gsα-mediated signaling following PTH treatment when compared to *WT*, suggest the presence of biallelic expression of *Gsα* within the osteoblast lineage.

## Discussion

Our data provide direct evidence of distinct changes in cortical and trabecular bone architecture among adult *Gnas E1+/−* mice dependent upon the inheritance pattern of their mutation. Specifically, mice with maternally derived (*Gnas E1+/−m*) mutations display elevated cortical and trabecular bone parameters when compared to WT mice. We identified a direct correlation between these increased cortical and trabecular bone parameters and enhanced osteoblast activity (as measured by MAR and serum P1NP) within *Gnas E1+/−m* mice. These observations directly correlate with our previous findings of normal to increased BMD in PHP1A patients.^(42)^ Alternatively, mice with paternally derived (*Gnas E1+/−p*) mutations displayed evidence of reduced cortical bone parameters when compared to both *Gnas E1+/−m* and WT mice. Similarly, *Gnas E1+/−p* mice displayed evidence of normal to reduced osteoblast activity in conjunction with elevated osteoclast activity *in vivo*. These observations in *Gnas E1+/−p* mice directly correlate with the previous findings in the literature suggesting that impaired Gsα signaling leads to an osteopenic bone phenotype.^(36–41)^ These findings could have potential implications for patients with PPHP and have yet to be correlated.

Our observations of enhanced osteoblast activity within *Gnas E1+/−m* mice vary from previous findings performed by Ramaswamy *et al*. utilizing our AHO mouse model bred on a mixed 129SvEv × CD1 background.^(44)^ In this previous study, *Gnas E1+/−m* mice displayed a reduced MAR on the endocortical surface compared to littermate controls, whereas no significant differences were observed between *Gnas E1+/−p* and littermate controls.^(44)^ It is first important to note that this histomorphometric analysis was performed on mice younger than the time point that we have evaluated (this previous study’s analysis was performed at 2 weeks of age). Second, μCT analysis at 2 weeks revealed that both *Gnas E1+/−p* and *Gnas E1+/−m* mice displayed a reduced cortical thickness and cortical BV/TV when compared to littermate controls. This study performed additional μCT analysis at 3 months and displayed similar patterns in cortical bone parameters that we observed between *Gnas E1+/−p* and *Gnas E1+/−m* mice; however, no formal analysis of osteoblast number or function was performed past 2 weeks of age. Therefore, it is plausible that in the early phases of long bone development, osteoblast activity is impaired within *Gnas E1+/−m* mice. However, following the initial stages of bone modeling, osteoblast activity specifically increases within *Gnas E1+/−m* mice due to changes within the local bone microenvironment, whereas osteoblast activity remains normal to reduced in *Gnas E1+/−p* mice.

We also identified elevated rates of osteoclastogenesis and increased osteoclast size within both *Gnas E1+/−m* and *Gnas E1+/−p* primary BMM cultures when compared to WT. RT-PCR analysis of mature osteoclast cultures identified that both *Gnas E1+/−p* and *Gnas E1+/−m* cultures demonstrated significantly elevated mRNA expression of *Rank* and *Calcr* when compared to WT. These findings correlate with previous reports suggesting that *Gnas* serves as a negative regulator of osteoclastogenesis and osteoclast fusion, ^(37,44,50)^ and therefore, it is plausible that *Gnas* or adenylate cyclase activity may serve a role in directly modulating the surface expression of RANK receptors within preosteoclasts, leading to elevated rates of osteoclastogenesis.^(63,64)^

Despite similar observations within *Gnas E1+/−m* and *Gnas E1+/−p* osteoclasts cultures *in vitro*, only *Gnas E1+/−p* mice displayed evidence of statistically enhanced bone resorption activity when compared to WT *in vivo* as measured by histomorphometry and serum CTX-1 testing. *Gnas E1+/−m* mice displayed evidence of normal to slightly enhanced osteoclast activity compared to WT. We identified that both *Gnas E1+/−m* and *Gnas E1+/−p* mice displayed an overall reduction in total mRNA expression of both *Rankl* and *Opg* when compared to WT; however, *Gnas E1+/−p* mice displayed a significantly elevated *Rankl:Opg* mRNA expression ratio compared to WT due to a substantially greater reduction in *Opg* expression*. Gnas E1+/−m* mice maintained a *Rankl:Opg* ratio comparable to WT. This particular distinction was driven by the fact that *Gnas E1+/−m* displayed a more modest reduction in *Opg* mRNA expression when compared to *Gnas E1+/−p*. This relative difference in *Opg* expression provides an additional explanation into the distinct bone resorption findings among *Gnas E1+/−m* and *Gnas E1+/−p* mice *in vivo*. Additionally, given that OPG secretion is tightly regulated by osteoblast activity, it is possible that this distinction in bone resorptive activity may be secondary to an overall increase in osteoblast activity within *Gnas E1+/−m* mice.

When considering the imbalance within both osteoblast and osteoclast activity among *Gnas E1+/−m* and *Gnas E1+/−p* mice *in vivo*, we wanted to address whether these distinctions could be driven by inheritance-specific changes in Gsα mediated signaling within cell types involved in the bone remodeling process. Despite the observed changes in osteoblast activity *in vivo,* we observed no significant variations in *Gsα* expression, osteogenic differentiation or mineralization between *Gnas E1+/−m* and *Gnas E1+/−p* BMSCs when plated at equal cell densities *in vitro*. While we identified variations in *Alpl, Sp7* and *Bglap1* expression between *Gnas E1+/−m* and *Gnas E1+/−p* BMSCs, it is possible that these variations could have been the result of the heterogeneity of these cultures and the potential presence of contaminating osteoclasts within this system. Because of this, it is plausible that the transcriptional changes we observed could be reflective of potential variations in osteoblast-osteoclast interactions *in vitro.* In addition to observing no changes in osteogenic differentiation *in vitro,* both *Gnas E1+/−m* and *Gnas E1+/−p* mice similarly displayed an impaired bioactivity of Gsα-coupled receptors when primary BMSCs were treated with PTH 1-34 when compared to WT. These data correlate with previous findings that suggest evidence of *Gsα* biallelic expression within cells of the mesenchymal lineage and their differentiated states of adipocytes and chondrocytes.^(18,26)^

We also demonstrated that *Gnas E1+/−m* osteoclasts displayed impaired calcitonin receptor bioactivity when compared to both *Gnas E1+/−p* and WT cultures *in vitro*. We identified that this reduction in bioactivity was directly associated with impaired Gsα-mediated signaling, given that supplementation of *Gnas E1+/−m* cultures with forskolin resulted in comparable expression of *Crem* and *Ramp3* when compared to WT and *Gnas E1+/−p* cultures. Although we did not observe any variations in *Gsα* transcription between *Gnas E1+/−p* and *Gnas E1+/−m* BMM cultures *in vitro* (Fig 5C), these inheritance-specific variations in receptor activity, in conjunction with previous clinical reports of calcitonin-resistance in PHP1A, raise the possibility of partial *Gsα* imprinting within the mature osteoclast. While the isolation of purified populations of mature osteoclasts has been a continual challenge within the field, a cluster of reports have recently described the potential of isolating purified populations from a mixed primary BMM culture system by the use of fluorescent reporter transgenic mouse models and Fluorescence Activated Cell Sorting (FACS).^(65,66)^ The application of this emerging method in future studies is warranted, and these investigations could provide a more definitive insight into the prospect of differential *Gsα* expression within the mature osteoclast through genomic imprinting or through an unknown methylation-independent mechanism.

In addition to its role as an anti-resorptive pharmacologic target, the physiological role of the calcitonin receptor has gained interest, particularly with respect to its role in maintaining bone remodeling.^(67)^ This interest has stemmed from the paradoxical phenotype that was observed in multiple reports showing that mice with global inactivation of *Calcr* display an elevated BV/TV and bone formation rate without any changes in total osteoblast or osteoclast number.^(52,68)^ Additional studies of the lineage-specific deletion of *Calcr* revealed this anabolic phenotype *in vivo* was primarily driven by impaired calcitonin-receptor activity specifically within the osteoclast lineage (LysM-CRE) due to elevated sphingosine-1-phosphate (S1P) release.^(52)^ Lineage-specific loss of *Calcr* within the mesenchymal lineage (*Runx2-CRE*), however, resulted in no phenotypic variations in bone architecture, osteoblast or osteoclast activity, or changes in S1P.^(52)^ Interestingly, our data identified a significant reduction in *Calcr* mRNA expression in flushed tibia specimens as well as BMSC cultures within both *Gnas E1+/−m* and *Gnas E1+/−p* mice; however, only *Gnas E1+/−m* displayed impaired calcitonin receptor activity within the osteoclast lineage. *Gnas E1+/−m* osteoclasts also displayed a significantly elevated *Sphk1* expression when compared to *WT* cultures. Therefore, it is plausible that the observed distinctions in bone formation activity between *Gnas E1+/−m* and *Gnas E1+/−p* mice are driven by changes in sphingosine 1-phosphate (S1P) signaling by the osteoclast lineage due to calcitonin resistance. Although we did not observe any significant variations in plasma calcitonin measurements within *Gnas E1+/−m* mice (Supplemental Fig 2C), the identification of spontaneous hypercalcitoninemia within PHP1A is not typically observed clinically.^(45,47)^ Therefore, future provocative tests such as pentagastrin stimulation or salmon calcitonin administration are warranted in order to assess the potential of this calcitonin receptor-S1P signaling axis influencing bone remodeling within *Gnas E1+/−m* mice *in vivo*. However, given the complexity of AHO, particularly with respect to the potential of other hormones also influencing the bone microenvironment within *Gnas E1+/−m* and *Gnas E1+/−p* mice, an unbiased and comprehensive genome wide analysis is also warranted to more definitively address this question.

In conclusion, our data are the first evidence of distinctions in bone remodeling between *Gnas E1+/−m* and *Gnas E1+/−p* mice and raise the potential of partial calcitonin resistance specifically within osteoclasts in *Gnas E1+/−m* mice. We also provide evidence of distinctions in cortical and trabecular bone architecture that could directly impact patients with both PHP1A and PPHP and provides an etiology as to why we previously found that PHP1A patients display normal to increased bone mineral density.^(42)^ Further studies in patients with PPHP will be key in determining whether there are implications to overall bone health in terms of monitoring as well as treatment. Finally, these data also underscore that Gsα-signaling not only influences bone remodeling by influencing cellular specific activities but also serves an essential role in modulating cross-talk between mesenchymal- and myeloid-lineage cellular populations within the bone microenvironment.

## ACKNOWLEDGMENTS

The authors declare that they have no conflicts of interest. The authors are grateful to UConn Health’s Microtomography Core Facility (Doug Adams, Daniel Youngstrom and Renata Rydzik) and Cryohistology Core Facilities (Li Chen, Zhihua Wu, Xiaonan Xin) for their assistance in sample processing. In addition, the authors are grateful to Archana Sanjay for kindly providing access to oligonucleotides and her thoughtful review of our primary osteoclast cell culture phenotyping. This work was supported by Connecticut Children’s Albright Fund to E.L.G.-L. and National Institutes of Health Grants NIDCR T90DE021989-09 to P.M. and NICHD R21 HD078864 to E.L.G-L.

---

**Supplemental Figure 1:**
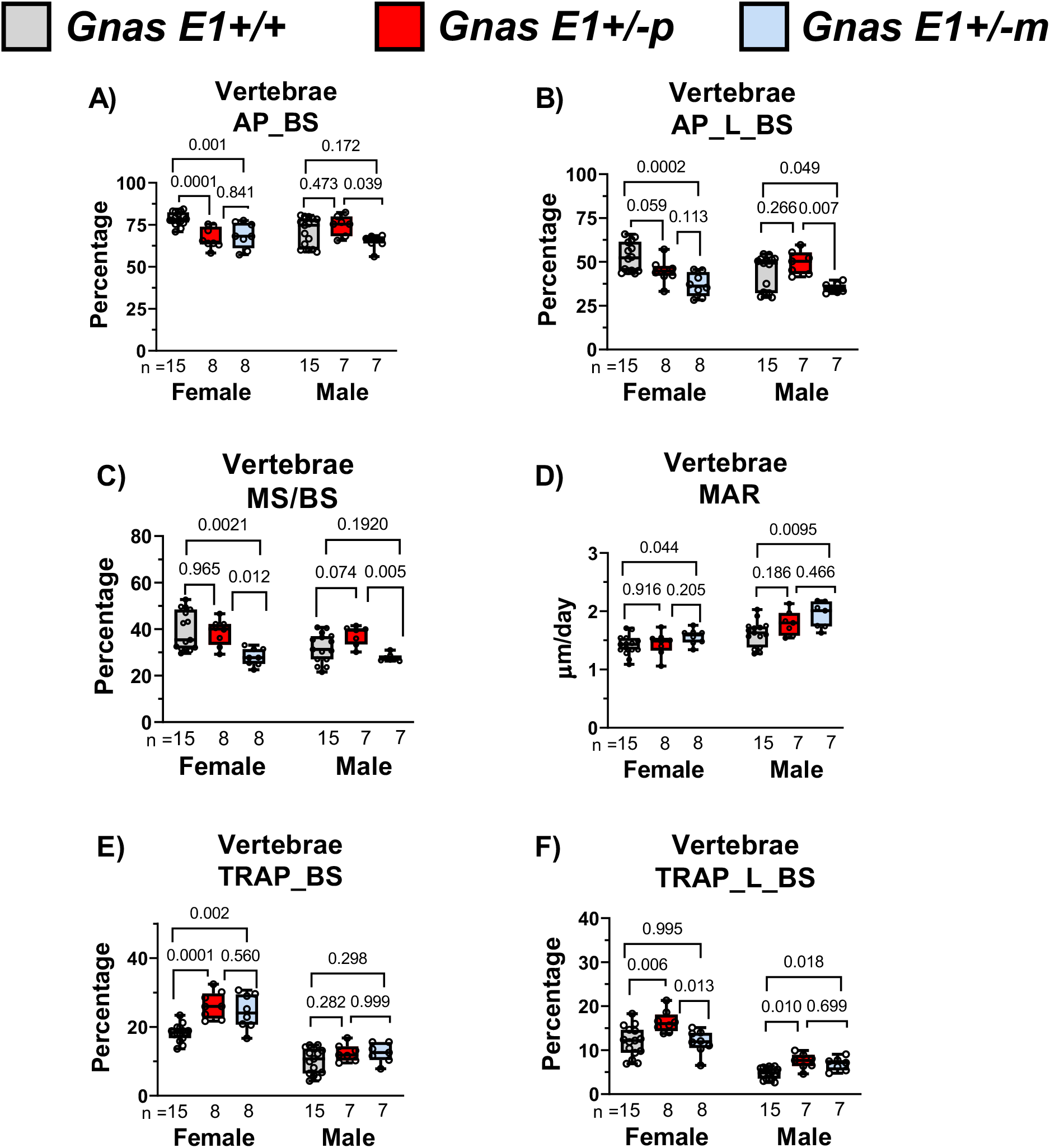
*Gnas* heterozygous inactivation differentially affects bone formation and resorption in the lumbar vertebrae. (A) Significant reductions in total alkaline phosphatase activity on the bone surface (AP_BS) was observed within female *Gnas E1+/−p* and *Gnas +/− m* lumbar vertebrae sections compared to WT. No significant differences were observed in male specimens across all genotypes. (B) *Gnas E1+/−m* mice displayed a reduction in the total number of actively mineralizing osteoblasts (AP_L_BS) when compared to WT and *Gnas E1+/−p* mice. (C) Dynamic histomorphometry on the vertebral trabecular surface revealed *Gnas E1+/−m* mice displayed a significant reduction in MS/BS when compared to both *WT* and *Gnas E1+/−p* mice (D) *Gnas E1+/−m* mice displayed a significantly elevated mineral apposition rate on the vertebral trabecular surface when compared to WT. (E) Female *Gnas E1+/−p* mice demonstrated a significant increase in total TRAP activity on the bone surface (TRAP_BS) and (F) active bone remodeling sites (TRAP_L_BS) within the vertebrae when compared to WT mice. No significant variations were observed between female *Gnas E1+/−m* and WT mice. No significant phenotype was observed within male mice. Sample size per genotype per experiment is listed on each bar graph. All statistical tests completed using ANOVA with post-hoc Tukey test for multiple comparisons, and p-values are displayed for each comparison.

**Supplemental Figure 2:**
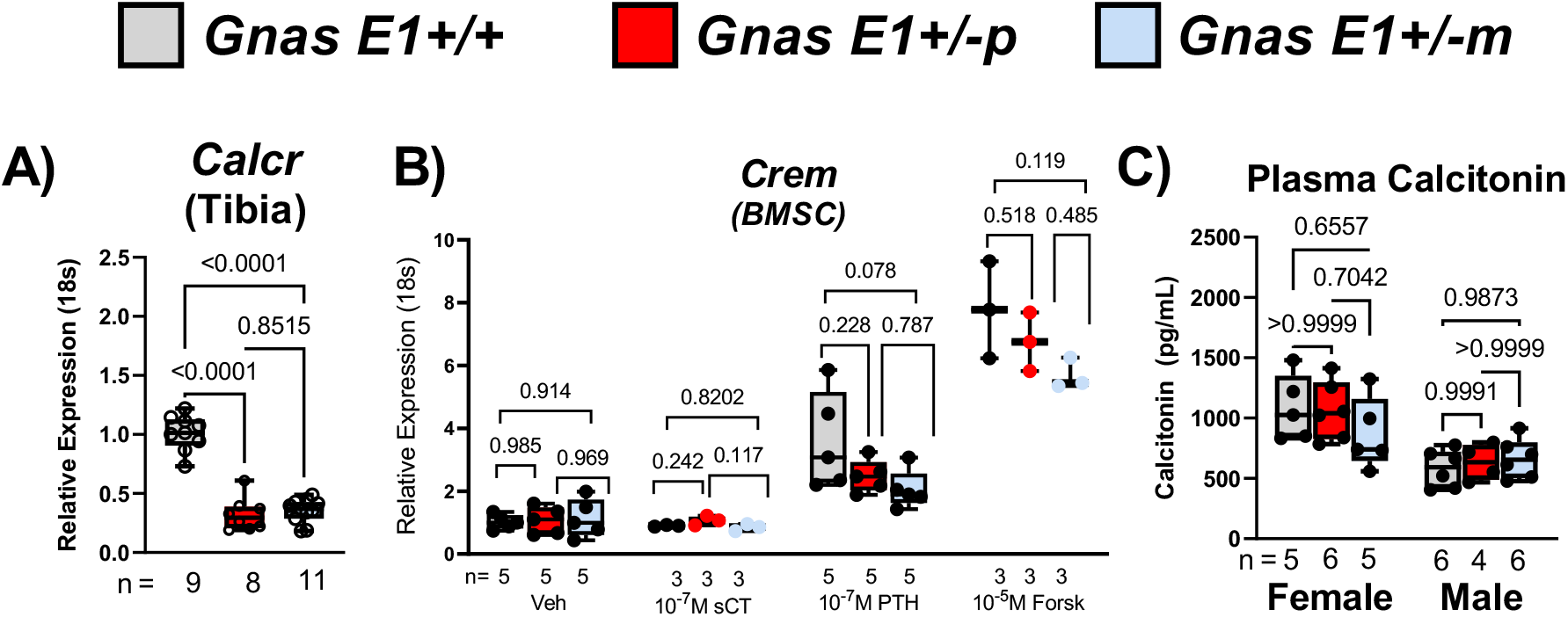
(A) RT-PCR analysis of flushed tibia diaphysis from male and female *Gnas E1+/−p* and *Gnas E1+/−m* mice displayed had a reduced mRNA expression of *Calcr* when compared to WT. (B) RT-PCR analysis of *Crem* mRNA expression of *Gnas E1+/−m, Gnas E1+/−p* and WT BMSCs following exposure to salmon CT, PTH, forskolin or vehicle controls for 6 hours. BMSCs overall displayed no significant response to sCT treatment. WT PTH-treated BMSCs displayed a significant increase in *Crem* when compared to vehicle controls, however, *Gnas E1+/−m* and *Gnas E1+/−p* BMSCs displayed no significant changes compared to vehicle controls. No significant variations were observed between PTH treated *WT* and *Gnas E1+/−m* or *Gnas E1+/−p* BMSCs.(C) Plasma calcitonin measurements obtained from 12 week old *WT* and *Gnas E1+/−* mice revealed no significant variations in calcitonin levels. Sample size per genotype per experiment is listed on each bar graph. All statistical tests in (A) and (C) completed using ANOVA with post-hoc Tukey test for multiple comparisons. Statistical analyses performed in (B) were completed using a two-way ANOVA with post-hoc Tukey test for multiple comparisons and p-values are displayed for each comparison.

